# The extent of protein hydration dictates the preference for heterogeneous or homogeneous nucleation generating either parallel or antiparallel β-sheet α-synuclein aggregates

**DOI:** 10.1101/2020.09.28.315325

**Authors:** José D. Camino, Pablo Gracia, Serene W. Chen, Jesús Sot, Igor de la Arada, Víctor Sebastián, José L. R. Arrondo, Félix M. Goñi, Christopher M. Dobson, Nunilo Cremades

## Abstract

α-Synuclein amyloid self-assembly is the hallmark of a number of neurodegenerative disorders, including Parkinson’s disease, although there is still very limited understanding about the factors and mechanisms that trigger this process. Primary nucleation has been observed to be initiated *in vitro* at hydrophobic/hydrophilic interfaces by heterogeneous nucleation generating parallel β-sheet aggregates, although no such interfaces have yet been identified *in vivo*. In this work, we have discovered that α-synuclein can self-assemble into amyloid aggregates by homogeneous nucleation, without the need of an active surface, and with a preference for an antiparallel β-sheet arrangement. This particular structure has been previously proposed to be distinctive of stable toxic oligomers and we here demonstrate that it indeed represents the most stable structure of the preferred amyloid pathway triggered by homogeneous nucleation under limited hydration conditions, including those encountered inside α-synuclein droplets generated by liquid-liquid phase separation. In addition, our results highlight the key role that water plays not only in modulating the transition free energy of amyloid nucleation, and thus governing the initiation of the process, but also in dictating the type of preferred primary nucleation and the type of amyloid polymorph generated depending on the extent of protein hydration. These findings are particularly relevant in the context of *in vivo* α-synuclein aggregation where the protein can encounter a variety of hydration conditions in different cellular microenvironments, including the vicinity of lipid membranes or the interior of membraneless compartments, which could lead to the formation of remarkably different amyloid polymorphs by either heterogeneous or homogeneous nucleation.

## Introduction

One of the key hallmarks of a variety of neurodegenerative disorders, collectively referred to as synucleinopathies, including Parkinson’s disease, is the aberrant aggregation of α-synuclein (αS)^1^, an intrinsically disordered protein highly abundant in neuronal synapses. αS aggregation can be reproduced *in vitro* and its kinetics are typically followed by means of thioflavin T (ThT) binding measurements, given the ability of this dye to bind specifically the cross-β structure of amyloid aggregates^2^. Using this type of assays, it has been established that amyloid aggregates are generally formed through nucleated self-assembly reactions, that follow typically a sigmoidal kinetics with a lag-phase (typically associated with a *t*_lag_) that in part reflects the energetically unfavorable primary nucleation step^3,4^, followed by a rapid-growth phase as a consequence of the more favorable processes of aggregate elongation, fragmentation, or secondary nucleation that take over once the early aggregates are formed, and a final stationary phase that reflects the consumption of monomers in the reaction and the equilibrium between monomers and aggregates^4^. While significant progress has been made towards understanding some of the microscopic processes that take place during amyloid self-assembly^4^, the processes that occur during primary nucleation, and the factors and mechanisms by which the process is initiated remain largely unknown. However, a detailed understanding of this process and its complex conformational landscape is a prerequisite for elucidating some of the molecular basis of pathology in Parkinson’s disease and related disorders, and for developing and evaluating effective disease-specific therapeutics to reduce or eliminate one of the underlying sources of toxicity in these diseases.

In *vitro*, αS amyloid aggregation has been typically triggered under strong stirring or shearing forces at physiological temperature, pH and ionic strength and ca. 50-100 µM protein concentrations^5,6^. Under these conditions, the air/water interface and/or the hydrophobic coatings of sample containers or stirring bars have been recently shown to play an essential role in triggering the initial self-assembly of αS through heterogeneous nucleation^6–8^, as a consequence of the particular amphipathic and surface-active properties of the protein^7^. Based on these findings, particular types of hydrophobic surfaces such as polytetrafluoroethylene (PTFE) beads, synthetic lipid vesicles or other surface-active materials^9–12^ have been recently used to accelerate primary nucleation of αS *in vitro*. Among all these experimental conditions, those involving lipid vesicles could represent the most physiologically relevant ones, although at the same time, at a neutral pH, synthetic vesicles composed of physiologically relevant lipids have been reported to be unable to trigger αS aggregation^12^, and only vesicles of very high contents of specific acidic lipids (typically small unilamellar vesicles, SUVs, containing 100 % dimyristoylphosphatidylserine, DMPS), seem to be efficient in catalyzing αS aggregation^12,13^.

Our understanding of the mechanisms that initiate αS aggregation inside the cells are even more elusive, although particular, yet undefined, cellular microenvironments appear to be required for the accumulation of αS amyloid aggregates inside cells, as the sole overexpression of the protein seem to be unable to trigger the *de novo* nucleation of αS amyloid formation, and additional cellular insults are typically required^14^. Importantly, recent experimental evidences show that the αS amyloid structures that have been resolved at high resolution, in all cases generated *in vitro* by heterogeneous nucleation at the air/water interface^15–18^, differ remarkably from the amyloid structures obtained from patient brain extracts^19,20^, suggesting alternative *in vivo* amyloid pathways from those explored until now *in vitro*.

In this context, we set out to expand the conformational landscape of amyloid aggregation and to understand the role of interfacial effects on the primary nucleation of the protein under different conditions. Interestingly, we have discovered that αS self-assemble in the absence of an active hydrophobic/hydrophilic interface by homogeneous nucleation under conditions of limited protein hydration, and that, when the protein undergoes this process, there is a preference for an antiparallel β-sheet arrangement of the amyloid structure, in contrast to the typical parallel β-sheet orientation acquired during heterogeneous primary nucleation. Interestingly, we have observed that the liquid-to-solid transition of αS droplets generated *in vitro* by liquid-liquid phase separation (LLPS) is likely triggered by this mechanism of homogeneous nucleation, in agreement with the particularly low water activity of such environments. Our findings, therefore, directly show the multiplicity of nucleation mechanisms and aggregate polymorphs within the αS amyloid aggregation landscape, and the role that water plays in selecting one type of mechanism and polymorph over others, suggesting that different cellular microenvironments with different protein hydration properties would lead to the formation of remarkably different amyloid polymorphs.

## Results and discussion

### Commonly used *in vitro* αS aggregation conditions require hydrophobic/hydrophilic interfaces

Under the most typical *in vitro* conditions used to trigger αS aggregation (PBS buffer, ca. 150 mM ionic strength, pH 7.4, 37 °C), we observed the formation of αS amyloid fibrils with the typical sigmoidal kinetic traces, although high protein concentrations and strong stirring were required (Fig. 1A). In order to compare aggregation kinetics and avoid possible effects of surface material of different sample containers^11^ (see Fig. S1), all the aggregation kinetics data shown in this study were performed in hydrophilic PEG-ylated plates^21,22^, unless otherwise stated. Under these conditions, the aggregation kinetic curves obtained were highly variable (Fig. 1A), as previously reported^5,23,24^, likely as a consequence of the stochastic and mutable nature of the air/water interface area of the sample at conditions of high-speed stirring. Indeed, we have observed that the nature and curvature of the air/water interface is determinant for the induction of αS aggregation under these conditions, in agreement with previous reports^6^, since the presence of air bubbles rather than a flat air/water interface particularly accelerates nucleation even in quiescent conditions (Fig. S2). The amyloid aggregates formed in this way are long, unbranched twisted fibrils of 10 ± 1 nm in height, 620 ± 400 nm in length (n = 50) and a twisting periodic pitch of 66 ± 3 nm (n = 25) according to atomic force microscopy (AFM) analysis. In order to analyze the effect of the air/water interface under this and other experimental conditions, we used hydrophilic poly(methyl methacrylate) (PMMA) caps that adequately fit into the wells and remove the air headspace, thus preventing the formation of the air/water interface^8^. When the caps were used, no aggregation was observed (grey line in Fig. 1A), and when the caps were eventually removed then aggregation was observed as expected (Fig. S2). Under quiescent conditions (without stirring), however, αS was not observed to aggregate for more than 250 h/10 days (even at 500 µM, Fig. S2A), in agreement with previous studies that have reported that, under similar conditions, αS was unable to aggregate even after several months of incubation^24^.

**Fig. 1.**
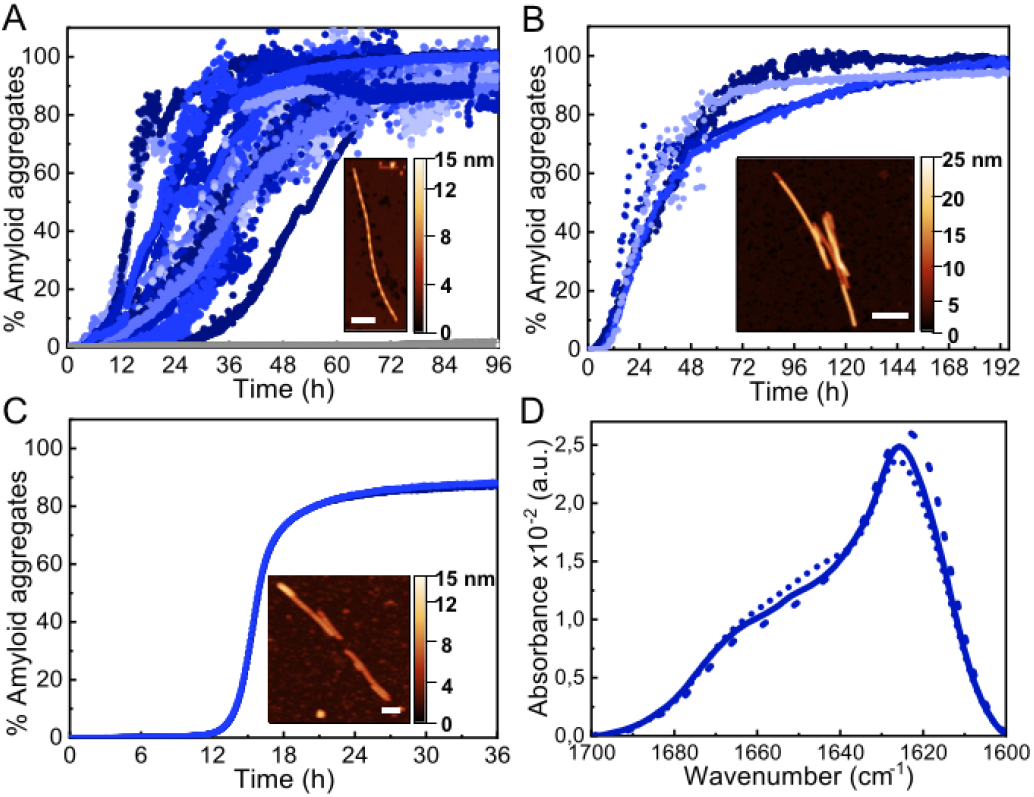
Characterization of αS aggregation at typical hydrophobic/hydrophilic interfaces. (A) Aggregation kinetics of 500 µM αS under shaking conditions (700 rpm). (B) Aggregation kinetics of 100 µM αS in quiescent conditions in the presence of a PTFE bead or (C) DMPS SUVs. The inset shows a representative AFM image of each type of aggregate. Scale bar: 200 nm. (D) IR spectra of the aggregates formed with shaking (solid line), or in the presence of a PTFE bead (dashed line) or SUVs (dotted line).

An alternative approach used to enhance *in vitro* αS aggregation involves the addition of PTFE beads, although this approach has been generally combined with agitation systems^25,26^. Here, we have analyzed the effect of adding a hydrophobic PTFE bead of a similar surface area to that of the air/water interface in our experiments (ca. 32 and 38 mm^2^, respectively) to the protein solution at quiescent conditions (100 µM αS in PBS buffer pH 7.4, 37 °C) and observed that nucleation at the surface of the PTFE bead was much faster than at the air/water interface (Fig. 1B), likely due to the stronger hydrophobic nature of PTFE as compared to air^27^. The amyloid fibrils formed under these conditions are typically 8.8 ± 0.8 nm in height and 360 ± 90 nm in length (n = 15), with no apparent twisted morphology according to AFM analysis. Synthetic lipid vesicles have also been used to trigger αS aggregation, although only SUVs composed of an artificially high content of saturated phosphatidylserine lipids (typically 100% DMPS) have been observed to trigger αS aggregation *in vitro* at nearly neutral pH^12,13^. We were able to reproduce the aggregation kinetics of αS under the above conditions (Fig. 1C). The fibrillar species so generated are typically 6.3 ± 0.6 nm in height and 800 ± 400 nm in length (n = 15) and lack the twisted morphology that was observed for the fibrils generated at the air/water interface under stirring. Despite the apparent differences in morphology, the amyloid fibrils generated at the three different types of hydrophobic/hydrophilic interfaces most typically used *in vitro* to trigger αS aggregation show remarkably similar infrared (IR) spectra (Fig. 1D) with the typical absorption band at ca. 1625-1615 cm^−1^ associated with intermolecular β-sheet amyloid aggregates^28,29^.

### Addition of low-to-mild alcohol concentrations accelerates αS amyloid nucleation

Biophysical characterization of protein folding and aggregation exploits often co-solvents to trigger changes in protein conformation or environment. Alcohols, particularly the fluorinated alcohol 2,2,2-trifluoroethanol (TFE), are perhaps the most used type of co-solvents to investigate protein aggregation and have been used in the past to study αS *in vitro* under shaking/stirring conditions^30,31^. Here we have used quiescent conditions and found that low concentrations of alcohols such as methanol (MeOH) or TFE (see Fig. S3 for the analysis of other types of alcohols), increased significantly the rate of αS aggregation (100 µM αS in PBS buffer pH 7.4, 37 °C, in quiescence; this conditions have been used for all further aggregation experiments) and increasing concentrations of alcohols resulted in shortened aggregation lag times (Fig. 2A, B). Although all the aggregates generated in the presence or absence of low concentrations of alcohols show remarkably similar IR spectra, with the typical amyloid-specific IR intense signal at ≈ 1625 cm^−1^ (Fig. 1D, 2C and S3C-D), significant variations in the morphology of the aggregates were observed by means of AFM imaging (Fig. 1A-C and 2D-G). For example, in the case of 5 % TFE, the aggregates show a fibrillar structure with an average of 10 ± 1 nm in height, and 600 ± 300 nm in length (n = 20) with a twisting periodic pitch of 61 ± 2 nm length, but the 10 % MeOH aggregates are preferentially globular, with average heights of 6 ± 2 nm and widths of 47 ± 8 nm (n = 50). Note that slight variations in the aggregation kinetics were observed when using other sample containers (Fig. S1), although the general effects and relative trends were very similar for the different conditions in all containers analyzed.

**Fig. 2.**
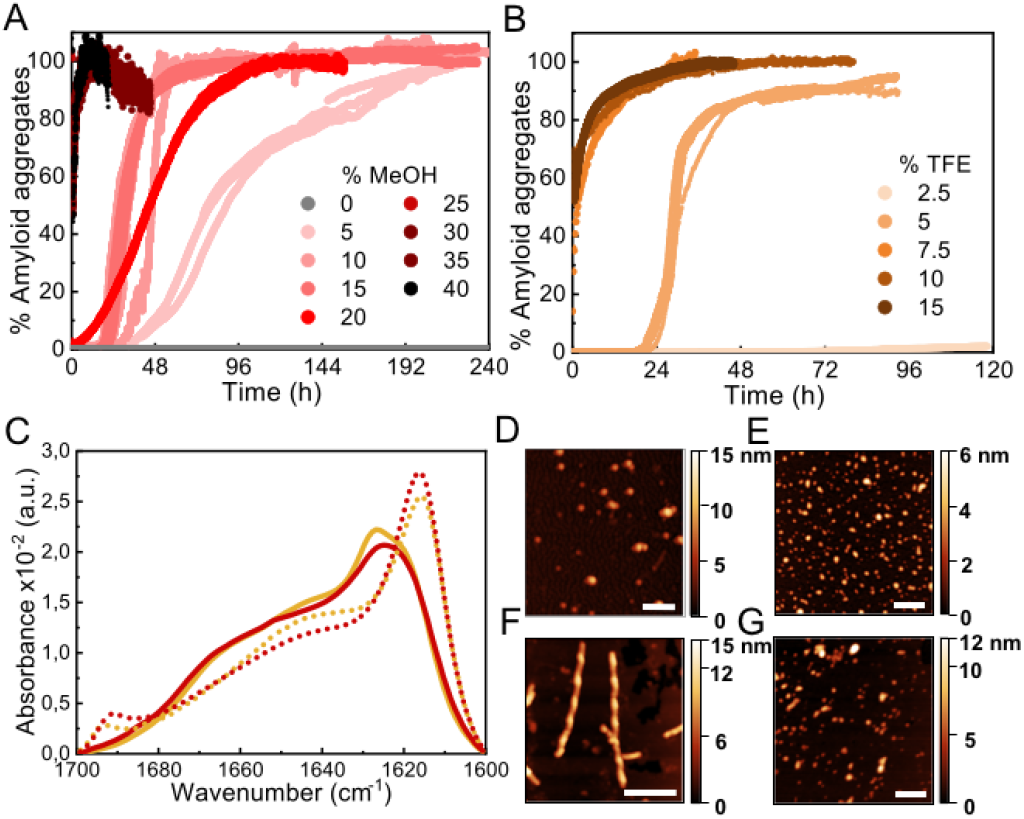
Characterization of αS aggregation in the presence of low-to-mild alcohol concentrations. (A-B) Aggregation kinetics of 100 µM αS in the presence of different concentrations of MeOH (A) or TFE (B). Three representative replicas are shown from more than 10 experiments with three different protein batches. (C) FT-IR spectra of aggregates formed in 10 % MeOH (red solid line), 35 % MeOH (red dotted line), 5 % TFE (orange solid line) or 15 % TFE (orange dotted line). (D-G) AFM images of aggregates formed in presence of 10 % MeOH (D), 35 % MeOH (E), 5 % TFE (F) and 15 % TFE (G). Scale bar: 200 nm.

Interestingly, we noticed that within a given alcohol concentration range, which varied depending on the type of alcohol used, but was surprisingly narrow in any case, there was a drastic change from a very long (typically >20) to a much shorter (<5-10 min) lag phase (Fig. 2 and S3). This phenomenon was observed, for example, when going from 5 to 7.5 % TFE or from 15 to 25 % MeOH. Under the fast nucleation conditions, the formed aggregates were preferentially globular, with average heights ≈ 5-6 nm and lengths ≈ 40-50 nm for the aggregates generated in 15 % TFE or 35 % MeOH, likely as a result of the preference for the stabilization of structures that maximize the volume of the aggregate while minimizing the surface area in contact to the poor hydrating solution. In all cases, however, the aggregates exhibited amyloid features including ThT binding (although the intensity of ThT fluorescence emission was significantly lowered as compared to the aggregates generated with longer lag phases), the amyloid IR fingerprint (intense signal corresponding to intermolecular β-sheet aggregates; Fig. 2C, and S3C-D), and the distinctive X-ray diffraction pattern indicative of amyloid cross-β structure (Fig. S4). In addition, these aggregates show seeding capabilities similar to the aggregates generated at lower alcohol concentrations, independently of their apparent size (Fig. S5), a feature characteristic of amyloid fibrillar-like aggregates. Note, however, that the IR spectra exhibit important peculiarities for the aggregates formed after short lag times (Fig. 2C, dotted lines), when compared to those requiring long lag times (Fig. 1D, Fig. 2C, continuous lines). Particularly, the main signal is shifted to ≈1615 cm^−1^, and a new, low-frequency signal appears at ≈1690 cm^−1^ (more on this below, in the context of the results in Fig. 6).

When we investigated the origin of the increased rates of αS aggregation by alcohols, we found that polarity alone cannot account for the results in the aggregation kinetics observed for the different types of alcohols analyzed (Fig. S6), in agreement with previous studies^29–31^. In addition to changes in the dielectric constant of the solution, alcohols have been reported to disrupt hydrophobic interactions and strongly affect the protein–water interactions, reducing the protein surface tension and consequently the protein hydration shell. Increasing concentrations of alcohols, particularly the hydrophobic ones such as TFE, lead, therefore, to increasing degrees of protein dehydration^32,33^. On the basis of this idea, Webb and colleagues proposed a de-solvation model assuming TFE-water mixtures as ideal solutions^31^, and the application of such model predicted that αS, and other proteins, would be preferentially de-solvated at moderated TFE concentrations in which they are prone to aggregate (although the nucleation occurring at interfaces was not considered in Webb’s model).

### Addition of high concentrations of salts, particularly kosmotropic salts, leads to accelerated αS amyloid nucleation

In order to investigate whether a decrease in protein hydration is a relevant factor in accelerating or even inducing αS aggregation, we analyzed the effect of salts, particularly kosmotropic salts, which have an opposing effect on electrostatics and hydrophobic interactions with respect to alcohols but, at high concentrations, are known to cause thinning of the hydration shell of proteins, similarly to alcohols, thus reducing protein solubility^34,35^. Upon increasing concentrations of salts in the 0.5-1 M range of ionic strength, both kosmotropic salts, such as NaCl and Na_2_SO_4_ (Fig. 3), and chaotropic salts, such as NaSCN (Fig. S7), a significant acceleration of αS amyloid aggregation was observed. At higher ionic strength concentrations,1-3 M for NaCl and 1-2 M for Na_2_SO_4_ or NaSCN, all salts used in this study behaved similarly, with aggregation kinetics that varied slightly with salt concentration, and with characteristic lag phases >10 h. However, there was a range of salt concentrations in which different behaviors between kosmotropic and chaotropic salts were observed. For the kosmotropic NaCl and Na_2_SO_4_ salts, the aggregation kinetics changed drastically at very high salt concentrations towards a mechanism with a remarkably fast nucleation step (with a lag-phase <5-10 min; Fig. 3A-B), while in the case of the highly chaotropic NaSCN salt, a sharp deceleration of the overall aggregation process was observed (Fig. S7). These results are in agreement with the ability of each type of salt used (Na_2_SO_4_, NaCl and NaSCN) to lower protein hydration and, thus to modify protein solubility (i.e. a strong kosmotropic, salting-out salt, a mild kosmotropic, salting out salt and a strong chaotropic. salting-in salt, respectively)^35^. Indeed, when analyzing in more detail the salt concentration range at which the slow-to-fast nucleation transition took place, we found that the preference for the fast nucleation mechanism occurred at lower salt concentrations for the highly kosmotropic Na_2_SO_4_ salt, as compared to the milder kosmotropic NaCl salt.

**Fig. 3.**
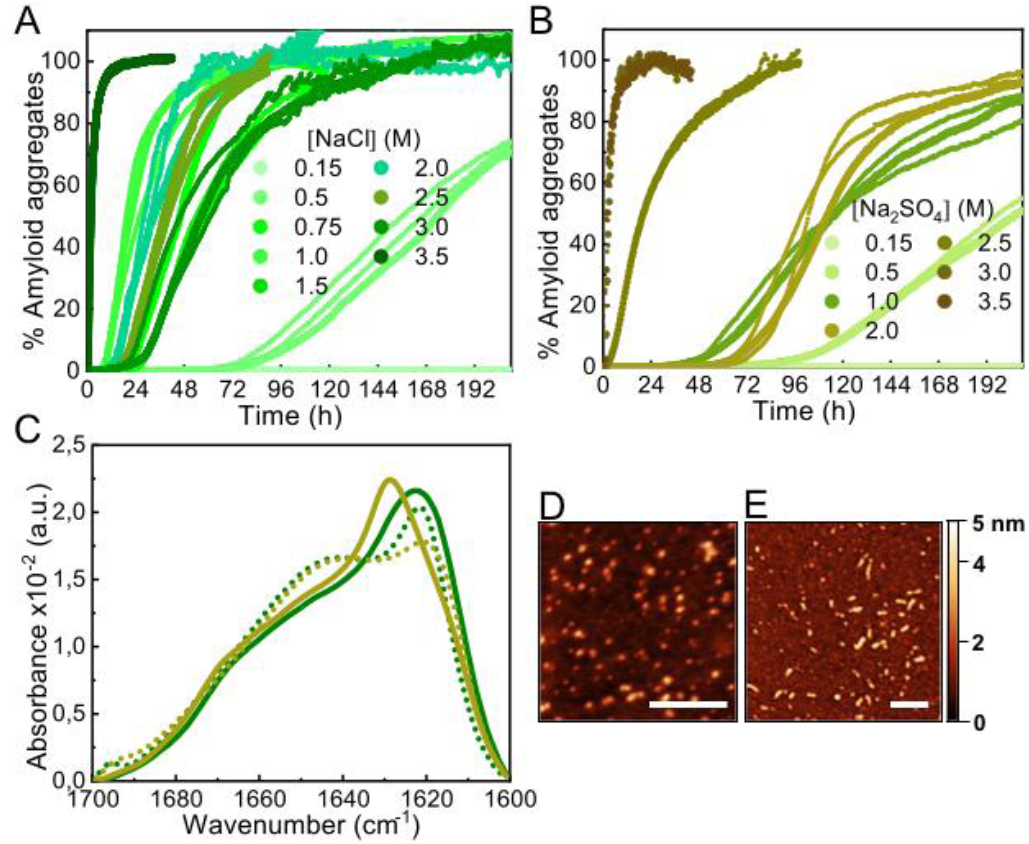
Characterization of αS aggregation in the presence of kosmotropic salts. (A-B) Aggregation kinetics of 100 µM αS incubated in the presence of different concentrations of NaCl (A) or Na_2_SO_4_ (B) (concentrations expressed in ionic strength). Three representative replicas are shown from more than 10 measurements with three different protein batches. (C) IR spectra of aggregates formed in 2 M NaCl (green solid line), 3.5 M NaCl (green dotted line), 2 M Na_2_SO_4_ (brown solid line) and 3 M Na_2_SO_4_ (brown dotted line) (concentrations expressed in ionic strength). Representative AFM images of aggregates formed in 2 M NaCl (D) and 3.5 M NaCl (E). Scale bar: 200 nm.

Also, as previously observed for low-to-mild concentrations of alcohols, the difference between the two αS aggregation kinetic regimes, with either long or remarkably short lag phases, was concomitant with a difference in the structural properties of the amyloid aggregates formed, as observed by IR analysis (Fig. 3C). The aggregates formed in high salt concentrations seemed to be preferentially small for all salts. For example the AFM-derived dimensions of the aggregates generated at 2 M or 3.5 M NaCl were 3.2 ± 0.2 nm in height, 20 ± 2 in length, *versus* 3.3 ± 0.2 nm in height, 23 ± 3 nm in length, respectively (n=50) (Fig. 3D, E).

### Addition of macromolecular crowders also accelerates αS nucleation

The third type of co-solvent or additive most frequently used for probing protein hydration changes is represented by inert synthetic polymers used as macromolecular crowding agents, which are preferentially excluded from the hydration shells of proteins, causing the preferential stabilization of compact protein conformations and complexes according to the excluded volume theory^36,37^, as well as a reduction in protein hydration^38,39^. In analogy with the results found with low alcohol concentrations and high salt concentrations, we found that mild concentrations (150 mg/ml) of three different types of crowding agents were able to accelerate αS aggregation (*t*_lag_ ≈ 10 h; Fig. 4A), in agreement with previous studies^30,40,41^, although in those studies strong protein sample agitation was used to trigger αS aggregation.

**Fig. 4.**
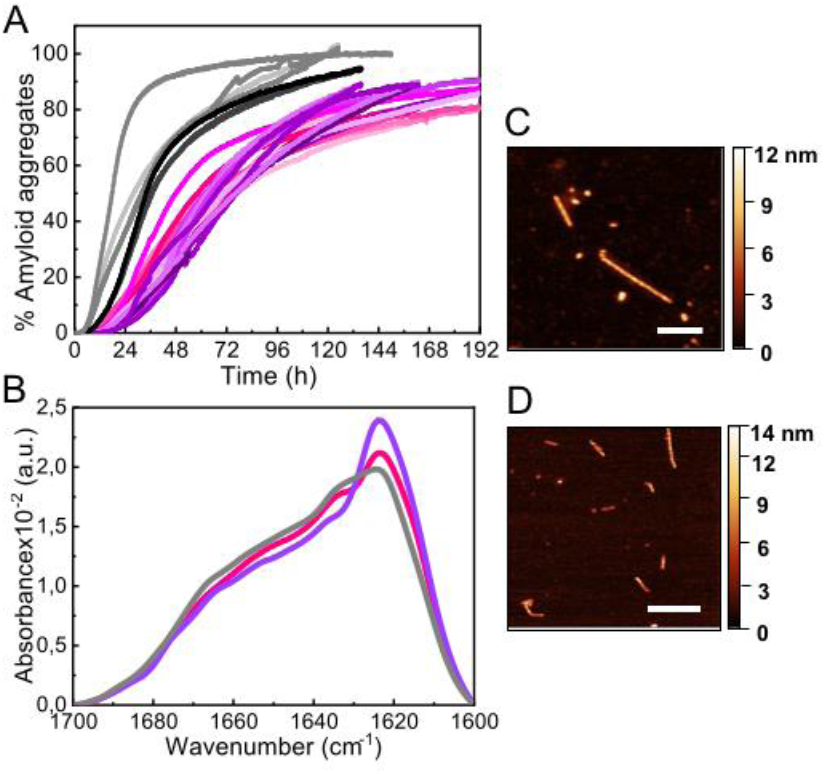
Characterization of αS aggregation in the presence of macromolecular crowders. (A) Aggregation kinetics of 100 µM αS incubated in the presence of 150 g/l dextran 70 (light-to-dark pink), ficoll 70 (light-to-dark purple) and PEG 8 (light-to-dark grey). Three representative replicas are shown from more than 10 measurements with three different protein batches. (B) IR spectra of aggregates formed in the presence of dextran 70 (pink), ficoll 70 (purple) and PEG 8 (grey). Representative AFM images of aggregates formed in dextran 70 (C) and ficoll 70 (D). Scale bar: 200 nm.

The aggregates formed under these conditions were preferentially fibrillar in the case of dextran 70 and ficoll 70 with similar dimensions according to AFM analysis: 6.0 ± 0.8 nm in height and 380 ± 170 nm in length (n = 30) in the case of dextran, and 6.3 ± 0.5 nm in height, 340 ± 170 nm in length (n = 15) for ficoll (Fig. 4C-D). In the case of PEG 8, however, more globular-like amyloid aggregates were observed (not shown). In any case, the three types of aggregates displayed an increase in ThT fluorescence signal and the β-sheet amyloid IR fingerprint. Indeed, we observed a remarkable similarity in the IR spectra of all the aggregates generated through slow nucleation process (see Fig. 1D, 2C, 3C and 4B). Upon increasing the crowding agent concentration up to 300-400 mg/ml, however, no change in amyloid aggregation mechanisms with a short lag phase was observed (not shown), unlike in the presence of alcohols or salts, probably because of the well-documented enhancement of surface-catalyzed protein fibril formation in the presence of macromolecular crowders^42^.

### αS forms amyloid aggregates without the presence of an active nucleation surface under limited hydration conditions

Given that at moderate alcohol concentrations and at salting-out concentrations of kosmotropic salts we observed a drastic acceleration in the apparent rate of αS nucleation (Fig. S9), we set out to characterize the type of primary nucleation that triggers the aggregation process at these conditions as compared to that occurring at lower additive concentrations. We first compared the kinetics of aggregation with and without hydrophilic caps designed to fit into the wells of the plates and remove the air/water interface^8^. These experiments were, however, rather complex and because of the slow aggregation reactions (lasting ca. 5-7 days) and the imperfect fit of the caps into the wells, the stochastic appearance of air bubbles in a significant number of samples was inevitable (Fig. S2B). For those samples where no bubbles were formed within the first 100h of incubation, no αS aggregation was observed under quiescent conditions in the presence of low concentrations of alcohols or salts (Fig. S10), indicating that under these conditions, αS heterogeneous nucleation at the air/water interface was accelerated with no signs of significant homogeneous nucleation.

At higher alcohol or salt concentrations, however, the nucleation process became remarkably fast, with identical behavior of the unmodified and the N-terminally acetylated forms of the protein (the latter being the most frequent physiological form in the cells^43^) (Fig. S11), thus alternative approaches were needed in order to analyze the role of the air/water interface on the amyloid nucleation process under these conditions. First, we performed surface tension experiments in a Langmuir tensiometer by the Wilhelmy method^44^ in order to analyze the ability of αS to adsorb and accumulate at the air/water interface under different conditions of either slow or fast primary nucleation. As expected, in the absence of MeOH, the addition of αS to the solution led to the partitioning of the protein between the bulk of the solution and the air/water interface, and the adsorption of the protein to the interface resulted in a significant reduction of surface tension that could be directly monitored in the Langmuir tensiometer (Fig. 5A). At protein concentrations above the saturation concentration in our experimental set up (ca. 60 nM of αS for an estimated air/water interface of 254.5 mm^2^), the protein generates a protein monolayer with a surface tension of 52 ± 2 mN/m (vs 70.2 ± 2 mN/m in the absence of protein) (see Fig. 5A, blue circles). When the same experiment was performed in the presence of 10 % MeOH, the interface saturation by αS was obtained at lower protein concentrations (≈25 nM), generating a protein monolayer with a constant surface tension of 52 ± 2 mN/m, indicating that the protein has a higher propensity to accumulate at the interface at 10 % MeOH, likely as a consequence of a higher propensity to acquire the amphipathic α-helical structure. Upon increasing MeOH concentration in the protein-free solution, the surface tension, however, decreased to ≈28 ± 2 mN/m at 35 % MeOH, already below the value of the surface tension generated by the αS monolayer (52 ± 2 mN/m), indicating that under these conditions αS would not partition into the air/water interface. Accordingly, we did not observe any variation in the surface tension of the 35 % MeOH solution upon addition of αS.

**Fig. 5.**
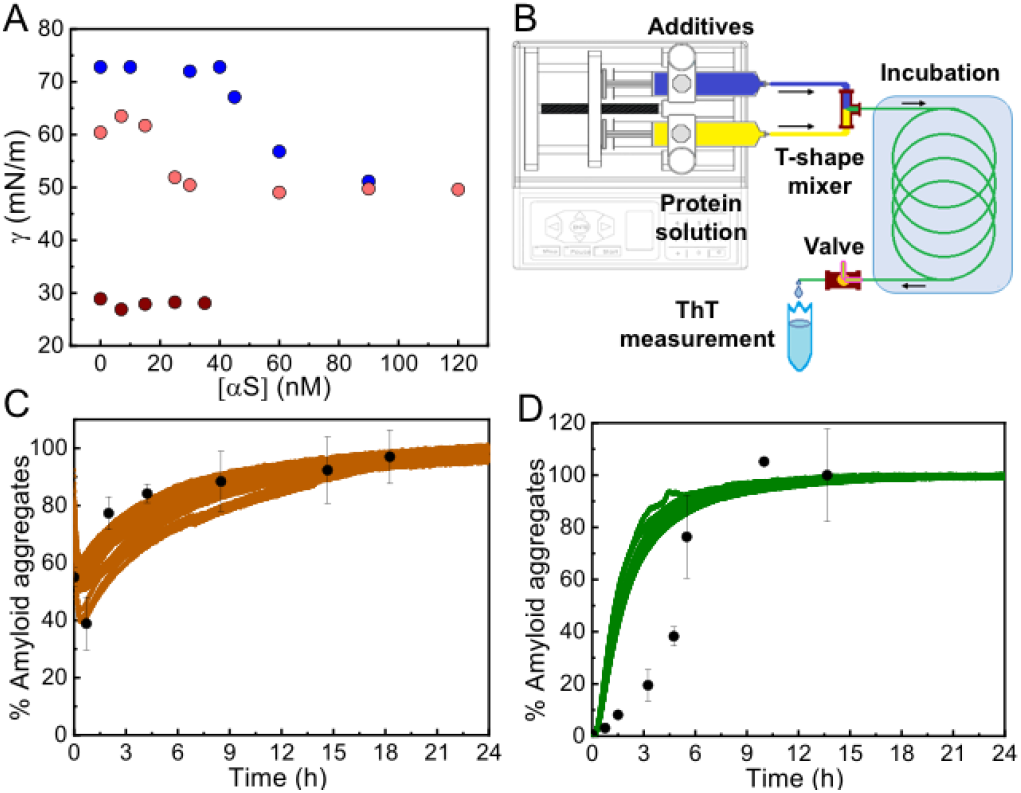
Propensity of αS to aggregate in the absence of an active hydrophobic/hydrophilic interface. (A) Surface tension measurements of αS solutions in PBS alone (blue), with 10 % MeOH (pink) or with 35 % MeOH (brown); average measurement error 0.82, 1.15 and 2.07 mN/m for each solution condition, respectively. (B) Outline of the microfluidic system used for the experiments shown in panels C-D. (C, D) Aggregation kinetics of 100 µM αS in the presence of 15 % TFE (C) or 3.5 M NaCl (D) incubated in the microfluidic system (black points). Bars represent standard errors for n=3-4 measurements per aggregation time point. Solid colored lines show data from standard aggregations in the microplate.

**Fig. 6.**
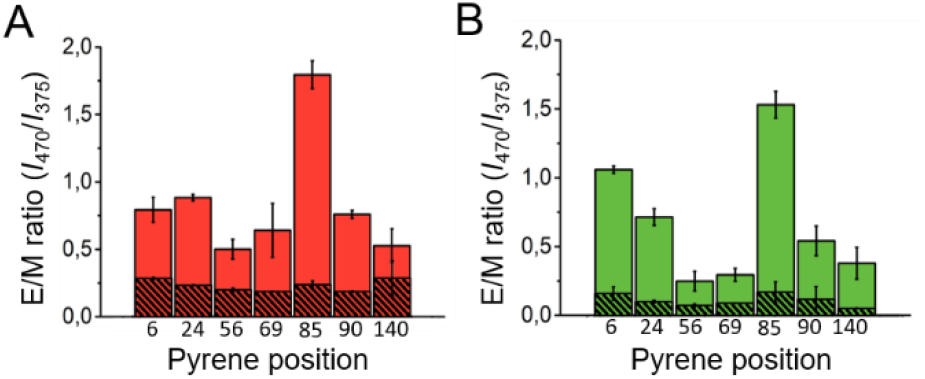
Intermolecular pyrene excimer formation analysis of the different β-sheet αS amyloid aggregates. (A) Pyrene E/M intensity ratio values obtained for the αS aggregates labeled with a 1:10 pyrene-αS / unlabeled-αS ratio at different sequence positions formed in PBS with 10 % MeOH (A, red), PBS with 35 % MeOH (A, shaded red), PBS with 2 M NaCl (B, green) or PBS with 3.5 M NaCl (shaded green), pH 7.4.

To further prove that αS forms amyloid aggregates in the absence of any hydrophobic surface that could initiate heterogeneous αS nucleation under limited hydration conditions, we used a microfluidic setup (see scheme of the system in Fig. 5B) in which the protein solution was mixed with the appropriate concentration of additives and incubated for a specific time only in contact with hydrophilic material (polyetheretherketone, PEEK). Using this device, we were able to reproduce the kinetics of aggregation that we observed previously in the plates, but this time without the presence of the air/water interface or any other hydrophobic surface that could initiate αS heterogeneous nucleation (Fig. 5C-D) (note that no apparent pre-formed aggregation nuclei were observed in the protein batch samples, see Fig. S12). Indeed, remarkably similar kinetics and yields of aggregation were obtained with drastically different surface materials and surface-to-volume ratios of the containers (in the case of the PEEK tubing the surface-to-volume ratio is 5-times higher than in the microplate wells). The small deviations observed between kinetics of αS aggregation in the presence of 3.5 M NaCl in the plate and in the microfluidic system (Fig. 5D) are most likely related to an acceleration of protein heterogeneous nucleation at the air/water interface in the plate under these conditions, since, at 3.5 M NaCl, and unlike at 35 % MeOH, the protein is expected to partition into the air/water interface where heterogeneous nucleation is probably triggered at faster rates than the nucleation in the bulk due to an increased local protein concentration. Taken together, our results indicate that αS can form amyloid aggregates under conditions of limited hydration in bulk solution due to homogeneous nucleation.

### αS preferentially forms parallel β-sheet amyloid aggregates by heterogeneous nucleation and antiparallel β-sheet amyloid aggregates by homogeneous nucleation

When we compared the IR spectra of the aggregates generated through either heterogeneous or homogeneous primary nucleation, we observed at least two remarkable differences (Fig. 2C and 3C). First we observed an additional band in the amide I region, at ca. 1690 cm^−1^, in the aggregates generated through homogeneous nucleation, indicative of the presence of antiparallel β-sheet structure^45–47^. Also, we noticed a general shift in the peak of the amyloid-characteristic amide I band from ca. 1625 cm^−1^ for the aggregates generated through heterogeneous nucleation to lower wavenumbers (ca. 1615 cm^−1^) for the aggregates generated through homogeneous nucleation. We attribute this red-shift of the amyloid-characteristic amide I band to a higher flexibility and/or solvent exposition of the β-sheet core^47,48^ in this type of aggregates.

In order to further corroborate the parallel and antiparallel arrangement of the β-sheet architecture of the two structural classes of amyloid αS aggregates, we analyzed the intermolecular fluorescence excimer formation of pyrene molecules attached to different positions of the polypeptide chain, particularly at positions 6, 24, 56, 69, 85, 90 and 140 in both types of amyloid aggregates. According to previous detailed structural analysis on parallel β-sheet αS fibrils (all formed through heterogeneous nucleation at the air/water interface)^15–18^, positions 56, 69, 85 and 90 are located in the β-sheet amyloid core, position 24 is located close to the N-terminal part of the amyloid core and positions 6 and 140 are located at the very N-terminal and C-terminal ends of the protein sequence, respectively. For this structural group of aggregates, therefore, when labeled with pyrene, we observed, as expected, excimer formation (emission band at around 470 nm) at the positions of the amyloid core, in addition to the monomeric pyrene emission bands (375nm-395nm) (see Fig. S13). This behavior was better visualized when using the excimer to monomer intensity ratio (E/M), which indeed can be used as a pyrene-pyrene proximity indicator^49^. Still, for the positions away from the core we did observe significant excimer formation, implying that in a parallel β-sheet arrangement even residues located in regions distant from the amyloid core are likely in close proximity, with transient intermolecular interactions, due to the flexible nature of those regions. However, in the case of the amyloid aggregates generated under conditions of very fast nucleation (Fig. 6, striped red and green), no significant excimer formation was observed for any of the αS positions analyzed, in agreement with a relative antiparallel orientation of the αS monomers within this type of amyloid aggregate.

### Homogeneous primary nucleation leads to amyloid aggregates with different thermodynamic stability with respect to the aggregates generated by heterogeneous nucleation

Our results indicate that αS aggregation can be triggered by different nucleation mechanisms leading to distinct amyloid polymorphs with different internal protofilament structure depending on the extent of protein hydration. This is particularly clear when alcohols, such as TFE or MeOH, are used to decrease protein hydration. At low alcohol concentrations, the formation of parallel β-sheet aggregates is preferred, while at higher concentrations the antiparallel configuration is stabilized. In order to analyze the influence of the extent of protein hydration on the stability of the resulting αS aggregates, we increased MeOH concentration of the solution and compared the percentage of parallel and antiparallel aggregates formed (Fig. 7 and Fig. S7).

**Fig. 7.**
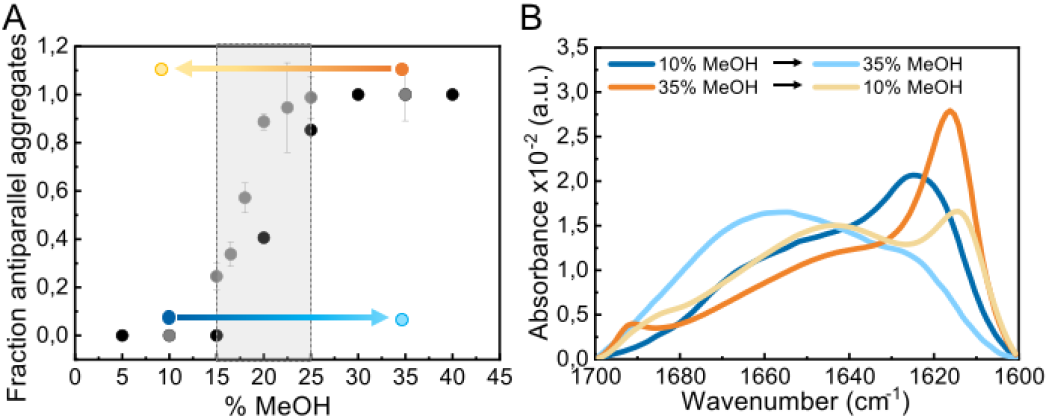
Impact of alcohol concentration on the aggregation behavior of αS. (A) Fraction of antiparallel aggregates found in the plateau phase of aggregation reactions of 100 µM αS in PBS pH 7.4 (37 °C) in the presence of different MeOH concentrations as derived from IR deconvolution (black dots) or pyrene E/M ratio (grey dots) analysis. For the pyrene data, error bars show the standard deviation of triplicate experiments, while for the IR data the uncertainty obtained from the global spectra fitting was 0.02 (see Supplementary Materials and Methods in Supplementary Information). The stability of parallel (generated at 10 % MeOH) and antiparallel (generated at 35 % MeOH) β-sheet aggregates under conditions at which the other type of aggregates is preferentially formed was analyzed (marked as blue and orange points and arrows, respectively). (B) Stability of the aggregates indicated in panel A by IR spectra.

As seen in Fig. 7A, the parallel β-sheet arrangement is preferentially formed at MeOH concentrations <15 %, while the antiparallel orientation is preferred at >25 % MeOH. Between 15-25 % MeOH, however, a mixture of both types of aggregates is observed, indicating that in this remarkably narrow regime of MeOH concentrations, thus, protein hydration conditions, both types of primary nucleation mechanisms are competing. When parallel β-sheet aggregates generated at low alcohol concentrations were transferred to conditions where the amyloid pathways of formation of antiparallel β-sheet aggregates were preferred and *vice versa* (Fig. 7A), the aggregates did not dissociate, according to SDS-PAGE analysis (not shown), as long as the MeOH concentration was higher than 5 %, and they maintained their original β-sheet arrangement (Fig. S13). Their structures, however, were severely destabilized (Fig. S13), with a significant increase in random coil content in both cases (Fig. 7B), which strongly affected their seeding capabilities (Fig. S5). Analogous results were obtained for parallel and antiparallel β-sheet aggregates generated with NaCl (Fig. S13). Note that when the antiparallel β-sheet aggregates were transferred to conditions of full protein hydration (PBS only), the aggregates fully dissociate very rapidly, while the parallel aggregates remained stable and reached a maximum β-sheet content (Fig. S14). These results indicate that the extent of protein hydration in different environmental conditions modulate the intermolecular interactions leading to a strong effect in both kinetics (in particular the early nucleation events) and thermodynamics of αS amyloid assembly.

### Antiparallel β-sheet aggregates represent the most stable structure of the preferred amyloid pathway triggered by homogeneous nucleation under limited hydration conditions

*In vitro*-generated amyloid aggregates of relevant disease-related proteins have long been observed to adopt in-register parallel β-sheet topology and the antiparallel β-sheet architecture has been, until now, assigned primarily for what has been referred to as stable (sometimes off-pathway) toxic oligomers^50,51^. Only very few cases of small synthetic peptides^52^ and disease-related amyloidogenic systems, such as the Iowa-mutant variant of Aβ peptide^53^, have been reported to form antiparallel β-sheet amyloid fibrils. We recently reported that a particularly toxic form of αS oligomers with an antiparallel β-sheet architecture was generated under conditions of very limited protein hydration, particularly during protein lyophilization^54^. In the present study, we have shown that αS can assemble into cross-β amyloid aggregates with significant antiparallel β-sheet structure in a range of limited hydration conditions and that these aggregates can have as much β-sheet content as the parallel β-sheet fibrils generated under more commonly used standard conditions according to IR spectra analysis (Table S1), despite their difference in the supramolecular structure. Indeed, the antiparallel β-sheet amyloid aggregates seem to have a smaller size as compared with the typical parallel β-sheet fibrils and, therefore, if they are eventually generated *in vivo*, they would be much more diffusive and, therefore, difficult to visualize or isolate.

The origin for the preference of the parallel or the antiparallel β-sheet structure in the αS amyloid aggregates is likely related with the type of primary nucleation under the particular solution conditions. When αS aggregation is triggered by heterogeneous nucleation, the pre-nucleus of amyloid structure formed at a given hydrophobic/hydrophilic interface would inevitably adopt a parallel intermolecular β-sheet arrangement given the restrictions in the disposition and orientation of the polypeptide chains anchored through their N-terminal amphipathic region to the interface. When the aggregation is triggered by homogeneous nucleation, however, there is no restriction in the orientation of the protein molecules in the bulk, and the antiparallel orientation of the β-sheets would be preferred over the parallel arrangement, as the stability of the hydrogen bonds in such configuration is generally higher^55,56^. An alternative explanation for the stabilization of the antiparallel arrangement at limited hydration conditions would be the strengthening of electrostatic interactions between the positively-charged N-terminal and the negatively-charged C-terminal regions of the protein. However, under conditions of high salt concentrations, where these interactions would be minimized, we still observe the formation of antiparallel β-sheet amyloid aggregates, and, importantly, the same behavior as the WT protein was observed for the C-terminally truncated αS variant (αS 1-103) (Fig. S11).

Our observation that limited hydration conditions promote the formation of αS amyloid aggregates rich in intermolecular antiparallel β-sheets has been, indeed, reported for other amyloidogenic peptides (derived from Aβ and Sup35) ^52,57^. In addition, a multitude of *a priori* non-amyloidogenic proteins belonging to different structural classes, as well as disordered peptides such as poly(L-lysine), have been reported to adopt an intermolecular antiparallel β-sheet conformation during the lyophilization process^58,59^ or under conditions of partial protein dehydration occurring at high temperatures^60^, or upon adsorption to hydrophobic surfaces without preferential protein orientations^61–63^. All these findings together, therefore, suggest that the formation of antiparallel β-sheet amyloid aggregates under limited hydration conditions may be a more general phenomenon of polypeptide chains, particularly when nucleating in bulk by homogeneous nucleation.

### The aggregates formed inside αS droplets generated by LLPS resemble those generated by homogeneous primary nucleation with a preference for an antiparallel β-sheet structure

Limited hydration environments can be found in the cell, particularly in the interior of protein-rich liquid droplets, also referred to as biomolecular condensates or membrane-less compartments, generated by protein-driven LLPS processes, and such environments have been reported to be particularly favorable for protein aggregation into amyloid structures (also referred to as liquid-to-solid transition) in a number of amyloidogenic proteins associated with disease^64,65^. Indeed, recent experimental evidences suggest a role of these protein droplets on the *in vivo* aggregation of these amyloidogenic proteins and the induction of pathology^66^. Very recently, it has been reported that αS droplet formation by LLPS precedes its aggregation in cellular^67^ and animal models^68^ and the liquid-to-solid transition of αS droplets has been recapitulated *in vitro*.

With the aim of elucidating if homogeneous primary nucleation and thus the formation of antiparallel β-sheet aggregates can be triggered inside αS droplets generated by LLPS, we have reproduced the conditions of reported *in vitro* αS droplet formation^68^ and observed aggregation inside the droplets very rapidly (within 2h of incubation, not shown) (Fig. 8B), with essentially all the protein being aggregated in less than 20 h (Fig. 8C), seen as ThT-positive aggregates. Under similar conditions without droplet formation, no aggregation was observed at the same time scale (Fig. S15). The rate of amyloid formation inside the droplets is, therefore, remarkably fast, already suggesting a homogeneous primary nucleation mechanism. When we investigated the nature of the amyloid aggregates formed, we observed that the aggregates exhibited relatively low ThT binding (not shown) and had relatively small sizes (2-h ultracentrifugation at 627.000 xg were required to obtain a pellet), features compatible with antiparallel β-sheet amyloid aggregates. Importantly, when we characterized the inter-molecular assembly of the β-strands in the aggregates using the pyrene excimer formation strategy, no excimer formation was observed when the pyrene probe was located in position 85, the position that shows the highest E/M ratio among all parallel arrangements characterized. Indeed, the E/M pattern (Fig. 8D) is identical to that of the previously characterized antiparallel β-sheet amyloid aggregates (Fig. 6).

**Fig. 8.**
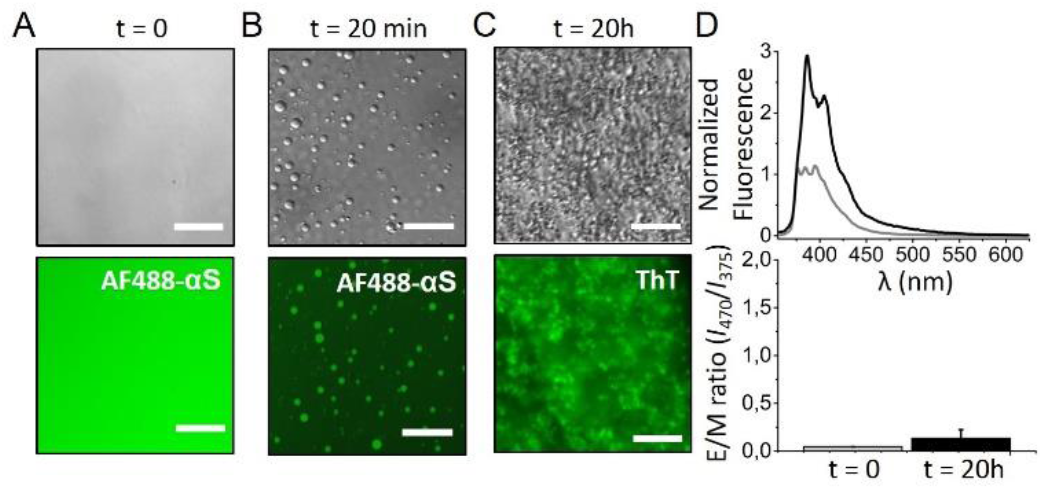
Analysis of the amyloid aggregates formed by the liquid-to-solid transition of αS droplets generated by LLPS. αS droplet formation was triggered by incubating 200 µM protein in 25 mM Tris, 50 mM NaCl, pH 7.4 in the presence of 10 % PEG8000 in a droplet set up (see Fig. S15). A) Representative images acquired immediately after sample preparation; Differential interference contrast (DIC, top panels) and widefield fluorescence microscopy (bottom panels). B) After 20 min of incubation, protein droplets were already observed. C) After 20 h of incubation, the protein sample was full of amyloid aggregates. For the fluorescence images in panels A and B, 1 µM AF488 αS was added to the protein solutions and the AF488 fluorescence signal was recorded. In the fluorescence image of panel C, bottom, all the protein was unlabeled and 100 µM ThT was added at time = 0. A GFP excitation/emission filter set was used for the fluorescence microscopy acquisitions in all cases. Scale bar: 25 µm. D) Representative pyrene spectra (top) and E/M ratio analysis (bottom) at time 0 h (grey) and 20 h (black) of incubation of a protein sample treated as in A-C) top panels, which also contained 20 µM pyrene-αS labeled at position 85. Error bars represent data obtained in two independent experiments.

Our data on the *in vitro* liquid-to-solid transition of αS droplets generated by LLPS strongly suggest, therefore, that the amyloid aggregates so generated have an antiparallel β-sheet structure, resembling the aggregates that we have observed to be generated by homogeneous primary nucleation in the bulk of the solution under poor hydration conditions in the presence of co-solvents or high concentrations of salts.

### Limited hydration conditions are required to trigger nucleation and the extent of protein hydration dictates the preferred type of primary nucleation and the type of amyloid polymorph generated

Despite the in principle intuitive relevance of protein hydration on the process of protein aggregation, the role of water in amyloid formation has been often overlooked. There is a limited number of studies in which this effect has been studied, most of them using a theoretical, molecular dynamics simulation approach^69–72^. In one of the few experimental studies using small amyloidogenic peptides, the authors used reverse micelles as a tool to modulate the accessible number of water molecules for the peptides, and demonstrated that the degree of hydration is an important determinant for the rate of initial self-assembly for the formation of both parallel and antiparallel β-sheet peptide aggregates^52^. We here have used different co-solvents and molecules whose only shared characteristic is that they are preferentially excluded from the protein surface, which results in the collapse of the protein hydration shell and, therefore, in a decreased protein hydration, thus promoting and stabilizing protein intra and intermolecular hydrogen bonding^73^. By systematically analyzing the effect of different concentrations of the different types of additives on the kinetics of αS aggregation and the structural types of aggregates formed under conditions where αS is unable to efficiently nucleate in a reasonable time, we have been able to observe a significant decrease in the lag phase of αS aggregation kinetics upon limited hydration conditions, with no apparent drastic changes in the structure or compaction of the monomeric structural ensemble of the protein at least under mild dehydrating conditions (Fig. S16). Importantly, two aggregation kinetic regimes were observed depending on the extent of protein hydration, one corresponding to kinetics with lag phases in the order of 20 h, i.e. at least two orders of magnitude shorter than in the absence of additives, and another regime with lag phases shorter than 5 min, i.e. at least four orders of magnitude shorter than in the absence of additives (see Fig. S9). Even if possible secondary processes could contribute to the reduction in the apparent lag phase under the different conditions^3^, our data show that under the conditions tested, a significant acceleration of primary nucleation occurs in the two additive concentration regimes with respect to the conditions in the absence of additives. Importantly, we here have demonstrated that while αS aggregation is triggered by heterogeneous primary nucleation in the first regime, homogeneous primary nucleation is the driving force of aggregation in the second regime, indicating that the extent of protein hydration is determinant for both types of primary nucleation. The initial self-assembly of amyloid structures either in the parallel or antiparallel arrangement, therefore, and in analogy to protein folding, requires overcoming a high de-solvation barrier^72^ arising from the removal of water molecules solvating the hydrophobic stretches that typically forms the amyloid core (the NAC region in the case of αS), as reported for the amyloid aggregation of the Aβ peptide^52,71^. Conditions of poor protein hydration, therefore, would significantly decrease the de-solvation barrier and the intermolecular hydrogen bonding barrier for nucleation.

At slightly reduced hydration conditions, αS nucleation can only take place effectively at hydrophobic/hydrophilic interfaces probably as a consequence of a local increase in protein concentration and a protein conformation/orientation selection at the interface, which, together with a more favorable intermolecular protein hydrogen bonding environment due to limited hydration conditions, allows for the formation of a sufficient number of simultaneous intermolecular interactions to offset the loss of polypeptide chains entropy, which is already low, as the protein molecules are anchored at the interface. αS, given its amphipathic nature, prompts a preferential adsorption at hydrophobic/hydrophilic interfaces in order to simultaneously maximize the hydrophilic interactions in the aqueous environment and the hydrophobic force at the hydrophobic surface^7^, and there, we and others have observed that the protein is able to nucleate the formation of amyloid aggregates^6,8^. Indeed, hydrophobic/hydrophilic interfaces have been found to be critical for the aggregation of many other amyloidogenic proteins and peptides that also share the amphiphilic character with αS, including disordered peptides such as amyloid β (Aβ peptide^74^ and folded proteins such as insulin^75^. Under more limited hydration conditions, however, we have observed that the nucleation process becomes favored, even at low micromolar protein concentrations (Fig. S17), and, therefore, it can take place very effectively in the bulk solution without the need of an active surface to concentrate the protein molecules or decrease their translational and configurational entropies. When homogeneous primary nucleation is then triggered, a different set of intermolecular interactions are established in the pre-nucleus which results in the formation a remarkably different amyloid aggregates with respect to those generated through heterogeneous nucleation, with a preference for an antiparallel β-sheet arrangement. At intermediate hydration conditions, as it would be expected, both homogeneous and heterogeneous nucleation were observed to co-exist, resulting in the formation of mixtures of aggregate polymorphs.

Interestingly, we have shown that primary homogeneous nucleation and the formation of antiparallel β-sheet amyloid aggregates could be a preferred amyloid pathway during the process of liquid-to-solid transition of αS droplets *in vivo*. Further investigation would be required to demonstrate the involvement of both homogeneous and heterogeneous nucleation in the formation of Lewy-body pathology and the induction of disease in *in vivo* models.

## Conclusions

In the present article, we have expanded the conformational landscape of amyloid aggregation and proved that αS is able to self-assemble into amyloid aggregates by homogeneous primary nucleation, without the need of an active surface, and that when the protein undergoes this process, there is a preference for an antiparallel β-sheet arrangement, in contrast to the parallel β-sheet architecture adopted when heterogeneous nucleation dominates. Indeed, the parallel arrangement is likely to be preferred in this case because of the restricted conformations that the protein is forced to adopt when anchored to a hydrophobic surface. Interestingly, the antiparallel intermolecular β-sheet structure has been previously observed in stable, particularly toxic oligomers of αS and other amyloidogenic systems and has been indeed proposed to be distinctive of these toxic species. We here demonstrate that the formation of such structure in αS represents an additional amyloid pathway to that previously explored, which is highly favoured under limited hydration conditions, with kinetics orders of magnitude faster than previously observed, and that it might indeed represent a general amyloid pathway of polypeptide chains when nucleating by homogeneous primary nucleation. Interestingly, we have observed that such nucleation mechanism is favored inside αS droplets generated *in vitro* by LLPS and might, therefore, be relevant in the *in vivo* liquid-to-solid transition of αS droplets, as well as, perhaps, that of other amyloidogenic proteins, an event proposed to be relevant in the pathology of amyloid-related neurodegenerative diseases^66^. In addition, we have found that the extent of protein hydration is a key determinant not only for triggering αS self-assembly (maintaining αS monomeric and preventing it from misfolding and self-assembly under highly hydrating conditions), but also for dictating the preference for the type of primary nucleation (heterogeneous vs homogeneous) and the type of structural amyloid polymorph generated (parallel vs antiparallel β-sheet structure). Because of the variety of hydration conditions in different cellular microenvironments, including some with very limited water accessibility such as the interior of phase separated protein rich microdroplets or membraneless compartments, our findings suggest that, *in vivo*, αS could undergo the formation of remarkably different amyloid polymorphs, by either heterogeneous or homogeneous nucleation, depending on the particular location of the protein in the cells, which could be ultimately related to the multiplicity of αS polymorphs that have been associated with distinct neurodegenerative disorders^76^. Interestingly, a decrease in the water content in brain cells, likely as a consequence of an increased total intracellular protein concentration with advancing age has been reported^77–79^, which may also contribute to the increased incidence of amyloid formation and, therefore, amyloid diseases in the aged population.

## Experimental

### Materials and methods

#### Protein expression and purification

Wild type (WT) αS as well as αS variants were expressed in Escherichia coli strain BL21 (DE3) and purified as described previously ^80,81^. In the case of the N-terminally acetylated variant, a simultaneous double transfection with αS plasmid and pNatB (pACYduet-naa20-naa25) plasmid (Addgene, UK) was used ^82^. The cysteine-containing αS variants were expressed and purified as described for the WT protein but including 5 mM DTT in all purification steps.

#### Aggregation assays

In ThT-monitored aggregation assays, 100 µM αS monomer was incubated in PBS buffer pH 7.4, 50 µM ThT, 0.01 % azide, in the presence of given concentrations co-solvents or salts at 37 °C until reaction was complete. 500 µM αS was used for aggregation in PBS in the absence of co-solvents or salts under shaking conditions (700 rpm using in situ orbital agitation in the plate reader). Non-Binding 96-Well Microplate (µClear®, Black, F-Bottom/Chimney Well) (Greiner bio-one North America Inc., USA) were used and the plates were covered with adhesive foil to prevent evaporation. All buffer samples and additive stock solutions were pre-filtered with 0.22 μm filters and both the multi-well plates and microfluidic devices were thoroughly cleaned before use. Moreover, some control experiments to exclude the presence of pre-formed nuclei in the protein batches were performed (Fig. S12). Kinetic reads were recorded in a FLUOstar plate reader (BMG Labtech, Germany); excitation at 450 ± 5 nm and emission at 485 ± 5 nm. See Supplementary Information for kinetic curves analysis. For pyrene-labeled αS, aggregation assays were performed as described for the WT protein with a 1:10 labeled-to-unlabeled αS ratio and containing 200 µM TCEP to prevent disulfide bridge formation between cysteines during the aggregation.

#### Fourier-Transform infrared (FT-IR) spectroscopy

αS end-point aggregates, after two centrifugation-resuspension cycles in order to remove unreacted monomers from the solution were resuspended in deuterated buffer to a final protein concentration of ca. 4 mg/ml. Samples were then deposited between two CaF_2_ polished windows separated by a PTFE Spacer (Harrick Scientific Products Inc., USA). Spectra were collected in transmission mode at room temperature using a VERTEX 70 FTIR Spectrometer (Bruker, USA) equipped with a cryogenic MCT detector cooled in liquid nitrogen. IR spectra were processed and analyzed using standard routines in OPUS (Bruker, USA), RAMOPN (NRC, National Research Council of Canada) and Spectra-Calc-Arithmetic© (Galactic Inc., USA)^45^. Global fitting analysis of IR spectra of αS aggregates generated at different MeOH concentrations were performed as indicated in Supp. Info.

#### Atomic force microscopy

Aggregated αS samples were diluted to a protein concentration 0.1-0.5 µM and deposited on cleaved Muscovite Mica V-5 (Electron Microscopy Sciences; Hatfield, Pensilvania, USA). Slides were washed with double distilled water and allowed to dry before imaging acquisition on a Bruker Multimode 8 (Bruker; Billerica, USA) using a FMG01 gold probe (NT-MDT Spectrum Instruments Ltd., Russia) in intermittent-contact mode in air. Images were processed using Gwyddion and the width measurements were corrected for the tip shape and size (10 nm).

#### Surface tension measurements

The surface tension of PBS solutions in the presence or absence of different MeOH and αS concentrations was measured using a DeltaPi 4-channel Langmuir tensiometer (Kibron Inc., Helsinki, Finland). Prior to measurements, the trough was thoroughly cleaned with ethanol and milli-Q water (18.2 MΩ cm) and dried with a vacuum pump. The probes were burned with a jet flame before and after the experiment. The tensiometer was calibrated with PBS at 25 °C under constant shaking and then MeOH was added when necessary at the indicated concentrations. The protein was added at the appropriate concentrations after buffer equilibration. The surface tension of the different solutions was recorded until an equilibrium value was reached (30 min - 2 h depending on the solution conditions) within the accuracy of the measurement (± 0.01 mN/m).

#### Generation of the microfluidic system

The microfluidic system was made of polyetheretherketone (PEEK), and was assembled by the following components: 1) Capillary tubing for additive inlets. The outer and inner diameters were 0.06 and 0.04 inches, respectively. 2) T-shape micromixer (IDEX Health and Science, USA) with a swept volume of 0.95 µl to promote an efficient and fast micromixing of additives. 3) Capillary tubing for protein incubation. The outer and inner diameters were 0.06 and 0.04 inches, respectively. The tubing length was 280 mm. 4) a 2-way valve to assure a tight air system shut-off. Reagents were injected in the microfluidic systems with a flow rate of 300 µl/min using a Harvard PHD Ultra® syringe pump. The surface-to-volume (S/V) ratio for the PEEK tubing where the protein sample was incubated was 3.93 (while a value of 0.77 was estimated for the wells of the PEG-ylated microplate).

## Acknowledgements

This work was supported by MINECO (RYC-2012-12068, NC), MINECO/FEDER, EU (BFU2015-64119-P, NC), MICIU/FEDER, EU (PGC2018-096335-B-100, NC), MICIU/FEDER, EU (RTI2018-099019-A-I00, VS), MCI/AEI/FEDER, UE (PGC2018-099857-B-I00, FMG) and Basque Government (IT1264-19, FMG and IT1196-19, JLRA), and Fundación Biofísica Bizkaia and the Basque Excellence Research Centre (BERC) program of the Basque Government (FMG, JLRA). We thank the Proteomics Unit of Servicios Científico Técnicos del CIBA (IACS-Universidad de Zaragoza), ProteoRed ISCIII member for mass spectrometry analysis and Advanced Microscopy Laboratory (LMA; Dr. José Luis Diez-Ferrer) for access to and help with AFM imaging. We also want to thank Dr. Ramón Hurtado for assistance in fiber X-ray diffraction, Dr. Janet Kumita for help with the acquisition of PMMA caps, Pablo F. Garrido for help with the FT-IR spectra global fitting and Dr. Pierpaolo Bruscolini and Prof. Paolo Arosio for insightful discussions.

