## Supplementary Material for "The extent of protein hydration dictates the preference for heterogeneous or homogeneous nucleation generating either parallel or antiparallel β-sheet α-synuclein aggregates"

---

a. *Biocomputation and Complex Systems Physics Institute (BIFI)-Joint Unit BIFI-IQFR (CSIC), University of Zaragoza, 50018 Zaragoza, Spain.*

b. *Centre for Misfolding Diseases, Department of Chemistry, University of Cambridge, Cambridge CB2 1EW, UK.*

c. *Biofisika Institute (CSIC, UPV/EHU), University of the Basque Country, Campus Universitario, B. Sarriena, 48940 Leioa, Spain.*

d. *Department of Chemical Engineering. Aragon Institute of Nanoscience (INA). Aragon Materials Science Institute, ICMA. Aragon Health Research Institute (IIS Aragon), University of Zaragoza, 50018 Zaragoza, Spain.*

e. *Networking Research Center on Bioengineering, Biomaterials and Nanomedicine, CIBER-BBN, 28029 Madrid, Spain.*

f. *Department of Biochemistry and Molecular Biology, University of the Basque Country, Campus Universitario, B. Sarriena, 48940 Leioa, Spain.*

† Deceased 8 September 2019. In memoriam.

### Supplementary Materials and Methods

**Kinetic data analysis.** Aggregation kinetic data were fitted to the following sigmoidal equation:

$$F = F_i + \frac{(F_f + m_f \cdot t)}{1 + e^{\frac{-(t-t_{50})}{\tau}}} \quad \text{[Eq. (1)]}$$

where $F$ is the fluorescence intensity, $F_i$ is the fluorescence signal at time zero, $F_f + m_f \cdot t$ describes the final baseline at the plateau phase, $t_{50}$ is the time at which $F$ is half of the maximum intensity and $\tau$ is a characteristic time constant that represents the inverse of the apparent growth rate. The lag time, $t_{lag}$ is estimated as $t_{50} - 2\tau$ as described previously[1]. This equation is commonly used to analyze amyloid formation kinetics without the assumption of any specific model, so it does not reflect the complexity underlying the amyloid aggregation process, but provides descriptive, empirical parameters that can be used in comparative studies.

**Determination of the aggregation yield.** The fraction of aggregated αS was estimated by quantifying the relative concentration of the insoluble and the soluble fractions. After reaction was complete (in the plateau phase), the sample was ultracentrifuged at 627.000 g for 90 minutes in a Beckman Coulter Optima® TLX (Beckman, USA) at room temperature, using a Beckman Coulter TLA 120.2 rotor. The soluble fraction was collected and the protein concentration of the soluble fraction determined spectrophotometrically (by absorbance at 275 nm, using a molar extinction coefficient of 5600 $M^{-1}.cm^{-1}$) for samples in the absence of ThT. For samples containing ThT, an aliquot of the soluble fraction was loaded in a 15 % acrylamide gel, together with an aliquot of the initial protein sample before aggregating, and the concentration of protein in the soluble fraction of the aggregated samples was estimated by comparing band intensities, after Coomassie staining, using the ImageJ software (NIH Image).

**Lipid Vesicle Preparation.** Small unilamellar vesicles (SUV) were prepared from dimyristoyl phosphatidylserine (DMPS, Avanti Polar Lipids, USA) by sonication as described previously [2]. The concentration of lipid stocks and the final solution of SUVs in HEPES buffer pH 6.5 was estimated from the Fiske phosphorus assay [3]. In this way the protocol for the generation of SUVs with a 100 % efficiency vas validated. The same protocol then was used to generate SUVs in phosphate buffer pH 6.5.

**Aggregation assays with hydrophilic caps.** The air/water interface was removed from the samples in individual wells by manual insertion of self-manufactured caps [4], which are made of poly(methylmethacrylate) (PMMA). PMMA was chosen as material for the caps because it is a clear acrylic polymer, relatively hydrophilic, and it has been shown to be inefficient in promoting αS aggregation [5]. 150 µl sample was deposited in the wells where the air/water interface had to be suppressed. The PMMA caps, previously moistened with buffer, were then placed vertically on top of each well and allowed to enter smoothly. Afterwards, the caps were gently pushed until fully insertion of the caps into the wells. Part of the sample spilled out during insertion of the caps, leaving 110-120 µl underneath. To minimize sample loss due to evaporation and the appearance of bubbles due to imperfect sealing, the caps were sealed with methyl cyanoacrylate.

**Estimation of the effective protein and ionic strength concentration in solutions with macromolecular crowders.** The effective or corrected concentration of αS and ionic strength of the protein solution as a consequence of the reduction of volume fraction available to the solvent by the presence of 150 mg/ml of dextran 70 or ficoll 70 was estimated, as described previously [6], to be 110.8 µM and 166.2 mM, respectively, for a solution of 100 µM of αS and 150 mM NaCl; these differences can be neglected in αS aggregation kinetics analysis.

**Seeding aggregation assays.** αS aggregates were ultracentrifuged at room temperature as described above and pellets were resuspended in the desired buffer. The seed stock samples were then sonicated in a bath sonicator (Ultrasonic Cleaner-VWR, USA) for 5 min, and the seed concentrations were determined by absorbance after disaggregating the seeds into monomers with 4 M guanidinium chloride (absorbance at 275 nm, molar extinction coefficient of 5600 $M^{-1}.cm^{-1}$). The seeding reactions were carried out at 37 °C with 25 µM monomeric αS and 3.75 µM of seeds in non-Binding 96-Well Microplate (µClear®, Black, F-Bottom/Chimney Well) (Greiner bio-one North America Inc., USA). ThT kinetic reads were recorded in a FLUOstar plate reader (BMG Labtech, Germany); excitation at 450 ± 5 nm and emission at 485 ± 5 nm.

**Labeling of αS with N-(1-pyrenyl) maleimide.** The cysteine-containing αS variants were labeled with N-(1-pyrenyl) maleimide (Santa Cruz Biotechnology). A stock solution of the pyrene reagent was prepared in DMSO. The DTT present in each cysteine αS variant solution was replaced with 5 mM Tris(2-carboxyethyl)phosphine (TCEP, Sigma Aldric), exchanging buffer with a Sephadex G-25, PD-10 desalting column (GE Healthcare). The labelling reactions were performed with 100 µM protein solutions in 25 mM Tris-NaCl, 150 mM NaCl, TCEP 5 mM, pH 7.25 at 4 °C in the dark, with a 5-fold molar excess of the pyrene reagent. The final DMSO concentration in the labelling reactions was, in all cases, lower than 1 %. The reactions were terminated at ca. 12-15 h by adding 10 mM DTT to the solutions, and the labeled protein was separated from the unreacted reagent with a PD-10 column. The labeling efficiency (10-95 %) was calculated by MALDI-TOF and, additionally, spectrophotometrically using a molar extinction coefficient of 36,000 $M^{-1}.cm^{-1}$ at 343 nm for pyrene and a correction for the pyrene absorbance at 275 nm using the same dye concentration in DMSO.

**Pyrene-labeled αS aggregation assays.** For pyrene-labeled αS, aggregation assays were performed as described for the WT protein with the following modifications: a 1:10 labeled-to-unlabeled ratio cysteine-containing αS mixture was aggregated at a total 100 µM protein concentration in PBS pH 7.4 containing 200 µM TCEP to prevent disulfide bridge formation between cysteines during the reaction. When required, MeOH or NaCl were added to the mixture at a given final concentration. After aggregation, the residual monomer was removed by ultracentrifugation as described for the WT protein. Pellets were solubilised in 100 µL of the aggregation buffer, sonicated for 1 min in a sonication bath and pyrene spectra were immediately measured. Bath sonication ensured adequate dispersion of the aggregated samples, with no significant aggregate clustering, while avoiding disaggregation and any apparent structural changes in the aggregates.

**Steady-state pyrene fluorescence spectroscopy.** The emission spectra of the aggregated pyrene-labeled αS variants excited at 343 nm were collected at room temperature in a Cary Eclipse Fluorescence Spectrophotometer (Varian, Palo Alto, California, United States) with slit-widths of 5/5 nm. An averaging time of 100 ms was used. The fluorescence spectra of the final aggregates of each pyrene-labeled variant were normalised to the intensity at 375 nm ($I_{375}$) and subsequently analysed to quantify the E/M ratio by dividing $I_{470}/I_{375}$, as described elsewhere [7–9].

**αS aggregate stability assays.** Identical protocols for IR and pyrene fluorescence analysis of aggregate stability were used. The pyrene-labelled aggregates were generated with a 1:10 labeled-to-unlabeled αS ratio as explained above. Once the aggregates were generated and washed to remove unreacted monomers from the solution, a first spectrum was collected corresponding to the aggregates under the same conditions as they were generated. Then, the aggregates were ultracentrifuged, as described above, resuspended in a buffer of equal composition as the aggregation buffer but with a different alcohol or salt concentration, bath-sonicated and incubated for 20 h at 37 °C. After this time, possible monomeric protein generated from aggregate disaggregation was removed from the protein sample by ultracentrifugation. The pellet was then resuspended in the same buffer and the aggregate solution was later bath-sonicated to avoid aggregate clustering due to ultracentrifugation. Then, the IR or pyrene fluorescence emission spectrum were collected, representing the aggregate spectra at the new alcohol concentration.

**Estimation of the fraction of parallel and antiparallel β-sheet aggregates at the end of the aggregation reactions at different MeOH concentrations by FT-IR spectra global analysis.** A global fitting analysis of the IR spectra of αS aggregates generated at different MeOH concentrations in the range of 5-40 % MeOH was performed in order to estimate the fraction of parallel and antiparallel β-sheet aggregates present at the end of the αS aggregation reactions under the different solution conditions. For this, the following equation was used:

$$\mathrm{X_i = X_{\nearrow\nearrow} \cdot a_i \cdot (A + B \cdot [MeOH]) + X_{\nearrow\swarrow} \cdot (1 - a_i) \cdot (C + D \cdot [MeOH])} \quad \text{[Eq. (2)]}$$

where the observed absorbance value at each wavenumber, $\mathrm{X_i}$, is assumed to be a linear combination of the values of each structural aggregate (parallel, $\mathrm{X_{\nearrow\nearrow}}$, or antiparallel, $\mathrm{X_{\nearrow\swarrow}}$) and its population ($\mathrm{a_i}$, and $1 - \mathrm{a_i}$, respectively). Good fittings were obtained only if the signals of the parallel and antiparallel structures were assumed to vary linearly with the concentration of MeOH, an estimation that is in good agreement with the experimental data shown in Figure S12.

**Dynamic Light Scattering (DLS) measurements.** Estimations of the hydrodynamic radius of the αS species in the protein stock solutions under different conditions were taken with a DynaPro NanoStar (Wyatt, USA) equipped with a Peltier temperature control. The DLS measurements were performed at 25 °C at a fixed angle of 90 ° collecting 20 acquisitions using a 2 s acquisition time. Protein samples were prepared at a 25 µM concentration with filtered (0.22 µm cellulose acetate syringe filters) buffers. Data analysis was performed with Dynamics software (version 6.12.03). An average of 10 measurements were performed for the statistical size analysis.

**Far-Ultraviolet Circular Dichroism (Far-UV CD) measurements.** CD spectra were recorded on a Chirascan™ spectropolarimeter (AppliedPhotophysic, UK) equipped with a Peltier temperature controller. The far-UV CD spectra (190-250 nm) was collected using a 0.1 cm pathlength quartz cuvette at 25 °C with 50 nm/min scan speed, 1 nm data pitch. Each data point was averaged over 10 data acquisitions. The measurements were carried at protein monomer concentrations of 5 µM. The ellipticity data was later converted to mean residue ellipticity data.

**X-ray diffraction.** αS aggregates were ultracentrifuged at room temperature as described above and pellets were resuspended at a final concentration of at least 5 mg/ml. A small volume of the solution was placed between 2 glass capillaries sealed on the far ends and allowed to dry at RT for 2 to 3 days, as described previously[10]. X-ray diffraction patterns were collected at room temperature on the BL13-XALOC beamline at the ALBA Synchrotron (Cerdanyola del Vallès, Barcelona, Spain) at a wavelength of 0.97 Å. The distance to detector was 61.5 cm and the resolution was 3 Å.

**Liquid-liquid phase separation assays.** 200 µM αS was incubated at 25 °C for 20h in Tris 25 mM, NaCl 50 mM, pH 7.4 in the presence or absence of 10 % PEG8000. For widefield fluorescence microscopy, either 1 µM AF488-labelled N122C αS or 100 µM thioflavin-T were included. For pyrene fluorescence spectroscopy and E/M ratio analysis, 10 % of the total protein concentration was pyrene-labelled at position 85 (Pyr-A85C αS). LLPS was initiated by casting a 10 µL droplet inside a non-binding microplate well (Greiner bio-one North America Inc., USA) for 20 minutes, then the wells were sealed and the reaction was allowed to proceed for 20 h in a humidity chamber to avoid evaporation. Assays were also performed in the bulk exactly in the same manner as described above with the sole exception that microplate wells were filled with 150 µL of the reaction mixture. Two independent triplicate assays were performed. For pyrene fluorescence spectroscopy, end-point LLPS reactions were ultracentrifuged for 2 h at 627.000 xg in a Beckman Coulter Optima® TLX (Beckman, USA) at room temperature, using a Beckman Coulter TLA 120.2 rotor, then the pellet was solubilized in the original reaction conditions and pyrene spectra were acquired as described above. Two independent assays were performed.

**Differential interference contrast (DIC) and widefield fluorescence (WF) microscopy.** Images were acquired on a Leica Dmi8 inverted fluorescence microscope (Leica Microsystems, Germany) at room temperature. A halogen lamp or a mercury metal halide bulb EL6000 (for DIC and WF imaging, respectively) served as illumination sources. For WF microscopy, the light was focused on and collected from the sample using a 40x air objective lens (Leica Microsystems, Germany) and the excitation and emission light was filtered with a standard GFP filter set with bandpass filters of 460-500 nm and 512-542 nm for excitation and emission, respectively. For DIC microscopy, the same objective was used to collect the reflected light. Collected light was detected on a Leica DFC7000 CCD camera (Leica Microsystems, Germany). Exposure times were 50 ms for DIC microscopy imaging and 20 ms for WF microscopy imaging. Intensity was software-enhanced for the ThT images for comparative purposes due to the low degree of ThT binding to these aggregates. Images were analysed using ImageJ (NIH, USA).

### Supporting Figures

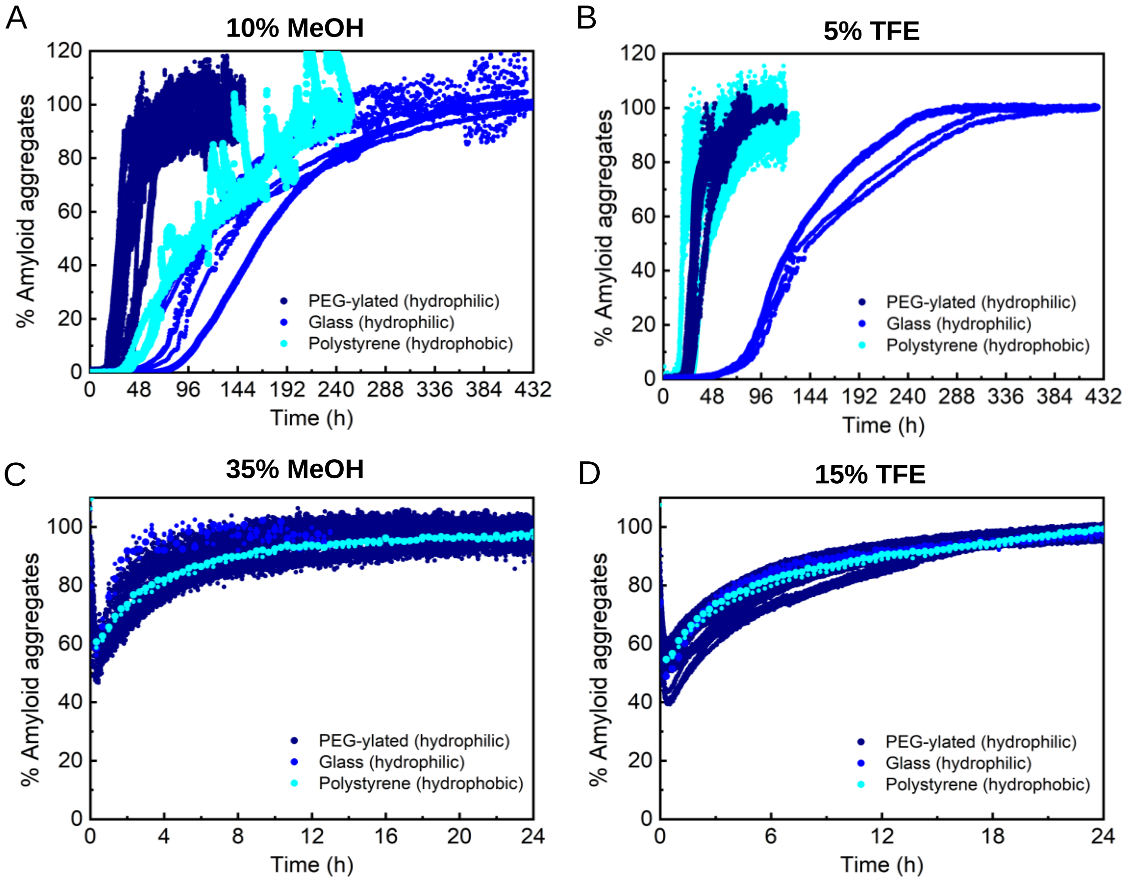


**Figure S1.** Impact of the container material on the aggregation kinetics of αS. Aggregation kinetics of 100 µM αS in PBS pH 7.4 at 37 °C in the presence of 50 µM ThT and 10 % MeOH (A), 5 % TFE (B), 35 % MeOH (C), or 15 % TFE (D). Each condition was assessed on PEG-ylated coated plates (dark blue), glass coated plates (standard blue), or naked polystryrene plates (cyan). While the aggregation kinetics vary significantly at conditions of slow nucleation such as 10 % MeOH or 5 % TFE when different sample container materials were used, no such differences were observed under conditions of fast nucleation such as 35 % MeOH or 15 % TFE, as expected for a heterogeneous nucleation- and a homogeneous nucleation-driven aggregation, respectively.

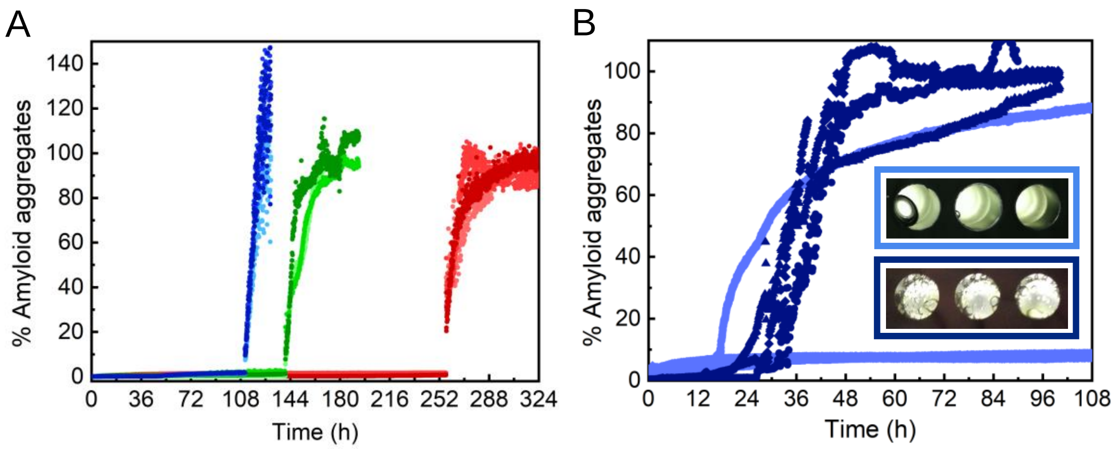


**Figure S2.** Impact of an air/water interface in the aggregation kinetics of αS. (A) Representative kinetic curves of 500 µM αS in PBS pH 7.4 (37 °C) in the presence of 50 µM ThT, incubated under shaking conditions (700 rpm) and with hydrophilic poly(methyl methacrylate) (PMMA) caps that fit tightly into the wells and remove the air headspace. For those samples where no air bubbles were formed, no aggregation was observed for long incubation periods. Eventually the caps were removed at different times and then aggregation was observed with the typical sigmoidal kinetics. (B) Representative kinetic curves of 100 µM αS in PBS pH 7.4 (37 °C) in the presence of 50 µM ThT, incubated under quiescent conditions and with PMMA caps. In some of the samples, and because the caps could not perfectly seal the wells, air bubbles appear during the kinetic process and aggregation is then triggered (the three replicas shown in dark blue and one of the three replicas shown in light blue, a picture of the corresponding wells for the two triplicates is shown in the inset). In two samples, there was no apparent formation of air bubbles and consequently no amyloid aggregation was observed (two of the three replicas shown in light blue). In the wells where air bubbles were formed, αS aggregation was triggered within the first two days of incubation despite no aggregation was observed for longer than 10-day incubations under the very same conditions (100 µM αS in PBS pH 7.4, 37 °C, in quiescence) without the PMMA caps.

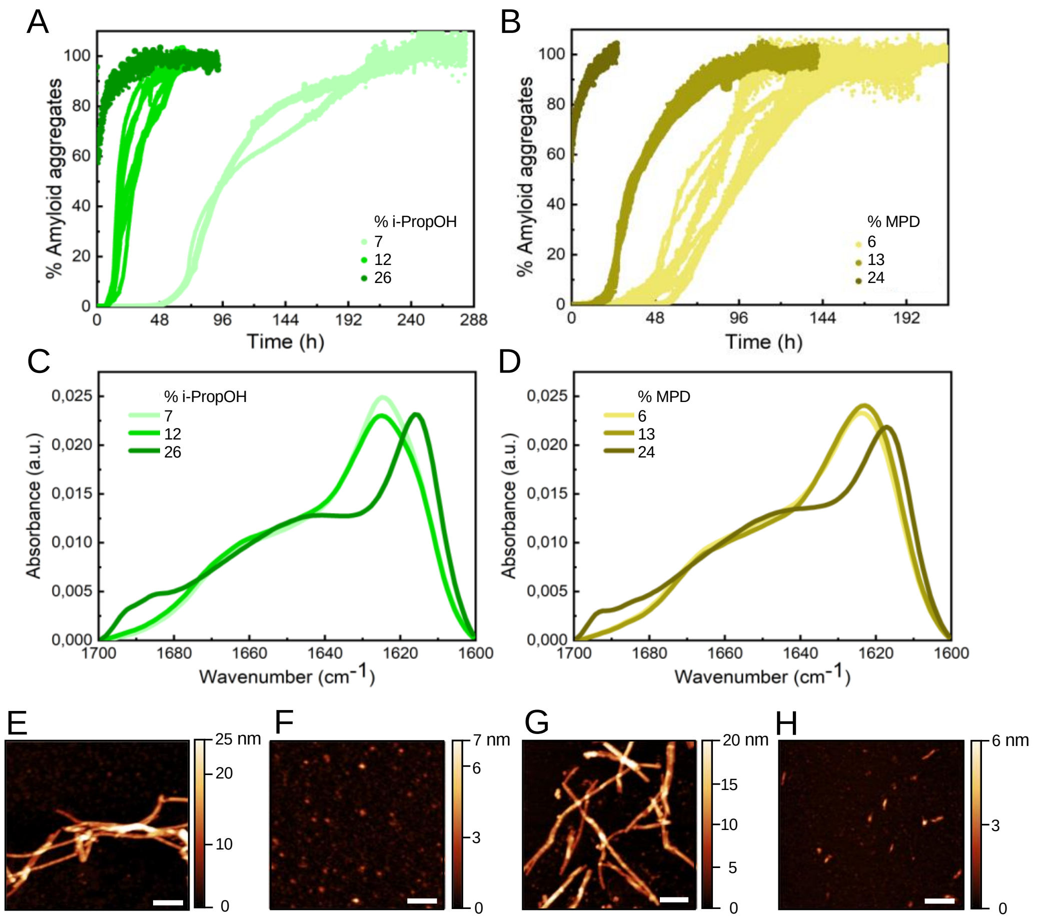


**Figure S3.** Characterization of αS aggregation in the presence of isopropanol (i-PropOH) or 2-methyl-2,4-pentanediol (MPD). Aggregation kinetics of 100 µM αS in PBS pH 7.4 (37 °C) in the presence of 50 µM ThT and different percentages of i-PropOH (A) or MPD (B). IR spectra of aggregates formed in the presence of i-PropOH (C) or MPD (D) (concentrations indicated in the figures) showing the two major different structural polymorphs of amyloid aggregates differing in their β-sheet rearrangement. Representative AFM images of the amyloid αS aggregates generated in the presence of 7 % (E) or 26 % (F) i-PropOH or in the presence of 6 % (G) or 24 % (H) MPD. Scale bar: 200 nm. In the presence of low alcohol concentrations, a preference for the formation of fibrillar aggregates was observed (for all the alcohols tested in this study except MeOH), while at higher concentrations, globular-like aggregates were more abundant.

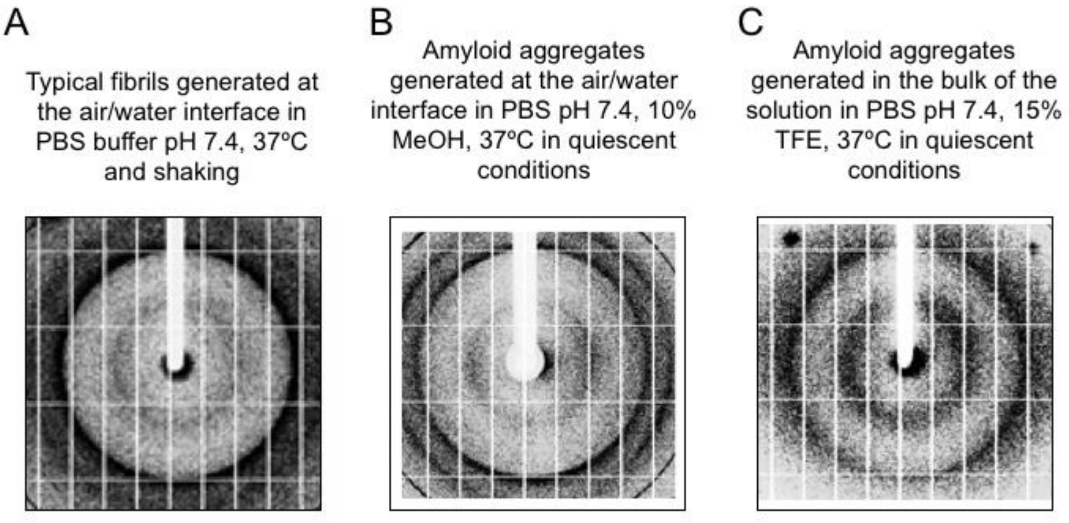


**Figure S4.** X-Ray diffraction pattern of αS amyloid aggregates. X-Ray diffraction pattern of the typical aggregates formed in PBS pH 7.4 (37 °C) at the air/water interface under shaking conditions (A) and the aggregates generated at 10 % MeOH (B) or with 15 % TFE (C) under quiescent conditions. In all cases, the X-ray diffraction patterns show the typical diffraction hallmark of the cross-β structure with the two characteristic distances that correspond to an inter-strand spacing in the aggregates of 4.68 ± 0.01 Å (A), 4.68 ± 0.01 Å (B) and 4.58 ± 0.01 Å (C) and an inter-sheet distance of 9.0 ± 0.1 Å (A), 9.0 ± 0.2 Å (B) and 9.3 ± 0.3 Å (C).

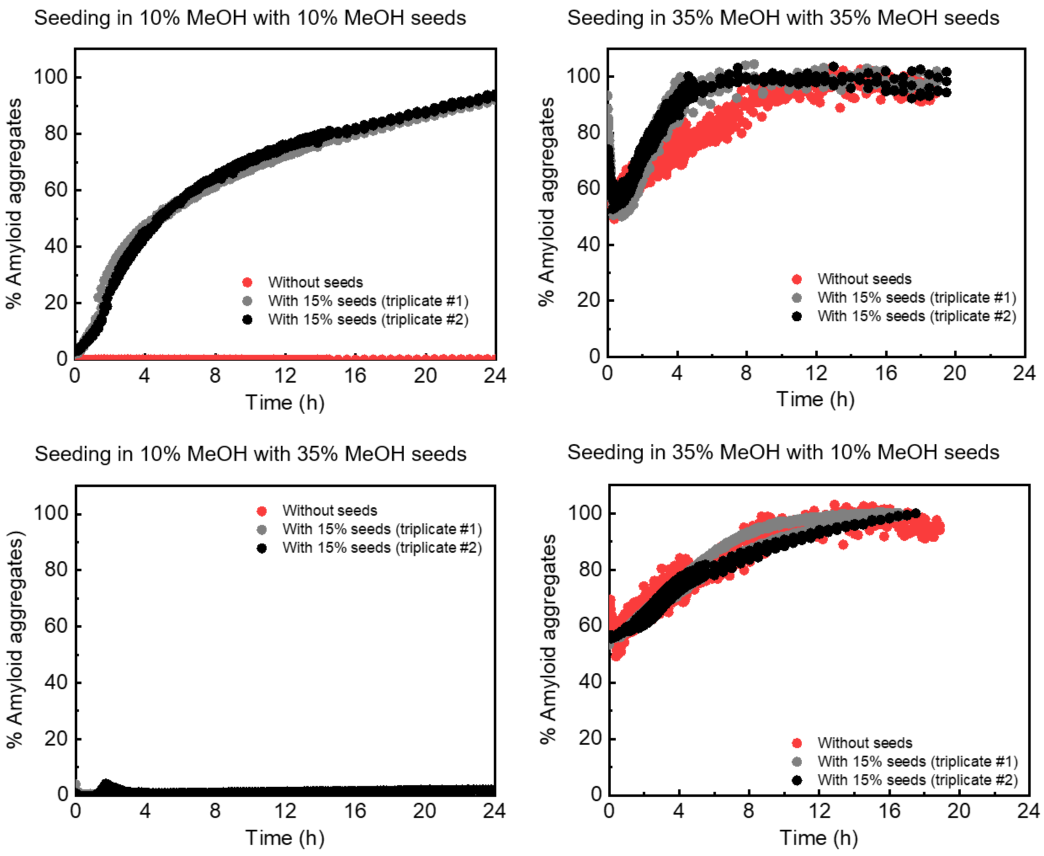


**Figure S5.** Seeding capabilities of the small amyloid aggregates generated under PBS, 10% MeOH or PBS, 35 % MeOH. Each type of aggregates was able to seed the formation of new amyloid aggregates when incubated with monomeric protein (25 µM monomeric protein, 3.75 µM seeds) under the conditions at which they were originally formed (top panels), demonstrating their amyloid fibrillar-like nature. However, the aggregates were not efficient at seeding under the conditions at which the other polymorph was more stable (bottom panels), in agreement with the strong destabilization of the aggregate structures upon solution transference as determined by FT-IR analysis (Fig. 7B in the main text).

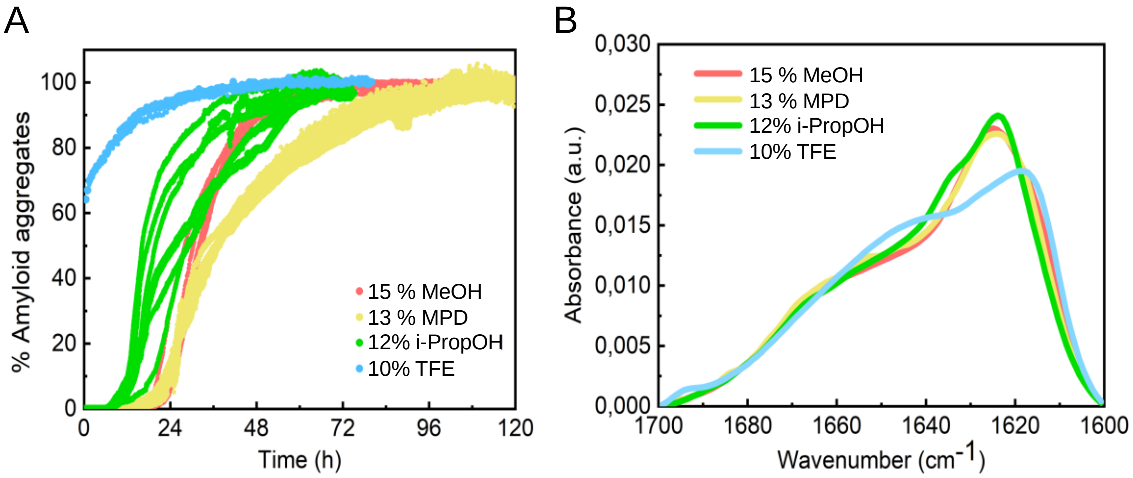


**Figure S6.** Kinetic and structural characterization of αS aggregates formed in the presence of different alcohols but with the same solution dielectric constant. Aggregation kinetics (A) and FT-IR spectra (B) of the aggregates generated with 100 μM αS in PBS pH 7.4 (37 °C) in the presence of 15 % MeOH (light red), 13 % MPD (yellow), 12 % i-PropOH (green) or 10 % TFE (cyan). The concentrations of the different alcohols were chosen to result in a solution with a dielectric constant equal or very close to 73.

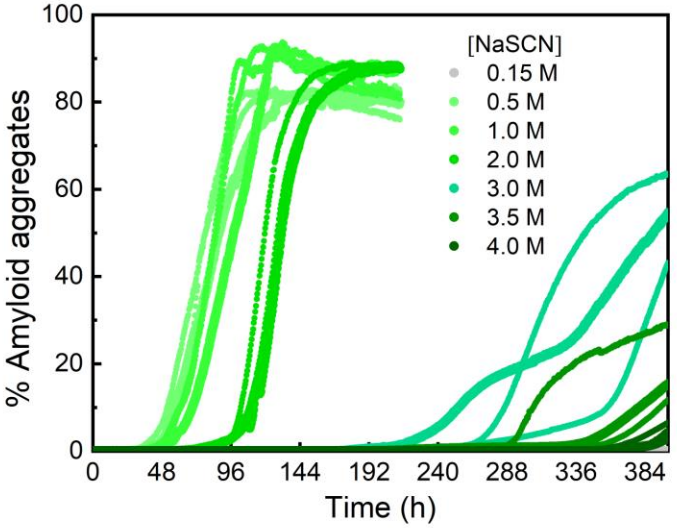


**Figure S7.** Kinetic characterization of αS aggregation formed in the presence of the highly chaotropic NaSCN salt. Aggregation kinetics of 100 µM αS in PBS pH 7.4 (37 °C) in the presence of 50 µM ThT and different concentrations of NaSCN. Upon a moderate increase in NaSCN concentration up to 2 M, there is a significant increase in the apparent rate of αS aggregation, in line with the effects observed for NaCl and $Na_2SO_4$. Above 2 M NaSCN, however, a drastic deceleration of the aggregation is observed, in stark contrast to the behaviour found for the kosmotropic salts (see Figure 3 in the main text).

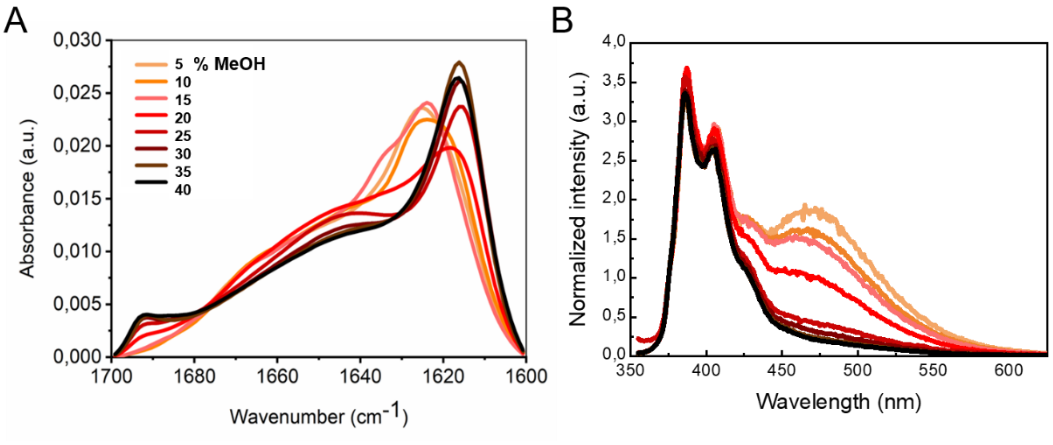


**Figure S8.** Structural analysis of αS aggregates formed in the presence of different percentages of MeOH. IR (A) and $I_{375}$-normalized pyrene fluorescence (B) spectra of aggregates formed with 100 μM αS in PBS pH 7.4 (37 °C) in the presence of different methanol concentrations. A clear transition from a parallel to an antiparallel β-sheet structure is observed using both methods, in both cases yielding virtually superimposable transition sigmoidal curves (see Figure 7A in main text).

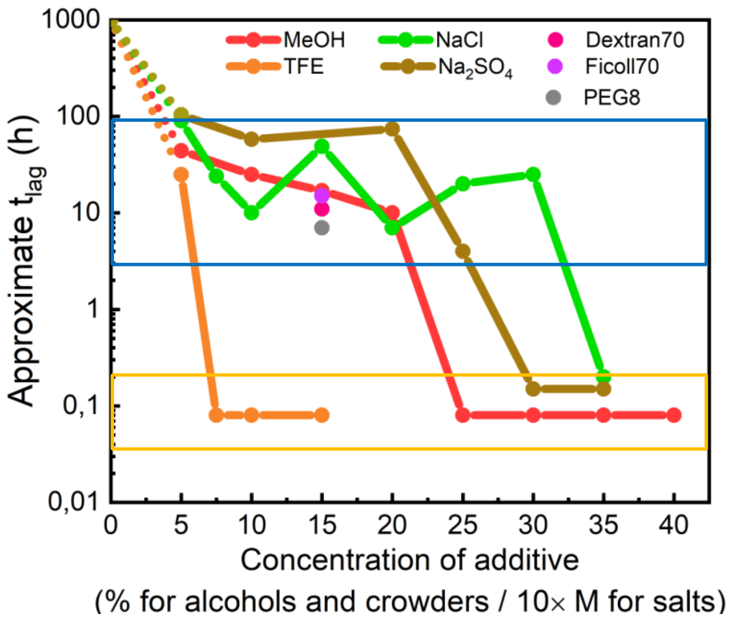


**Figure S9.** Analysis of the approximate duration of the αS aggregation lag phase with respect to the concentration of the different types of additives. In the absence of additives, no aggregation was observed under the conditions used in this study (100 µM αS in PBS, pH 7.4, at 37 °C, in quiescence) even after several months of incubation [11], so an arbitrary value of 1000 h (ca. 40 days) was chosen for plotting in order to visualize the acceleration of primary nucleation in the presence of the selected additives. In the presence of increasing concentrations of additives two main regimes in terms of duration of the lag phase were observed: one regime at which the lag phase of the aggregation reactions lasted typically longer than 10 h (conditions within the blue box) and another regime at which the lag phase was already finished after 5-10 min incubation (conditions within the yellow box), i.e. at least two orders of magnitude earlier than the typical lag phase of the previous regime. Interestingly, transition from one regime to the other is remarkably abrupt, suggesting that the aggregation reactions that fall within the two regimes are triggered by different primary nucleation mechanisms. The approximate $t_{lag}$ values for all the conditions, except the 0 value of concentration, were estimated from the analysis of the kinetic curves shown in Figure 2, 3 and 4 of the main text (see Supplementary Materials and Methods in Supplementary Information). The concentration of salts is given by ionic strength concentration.

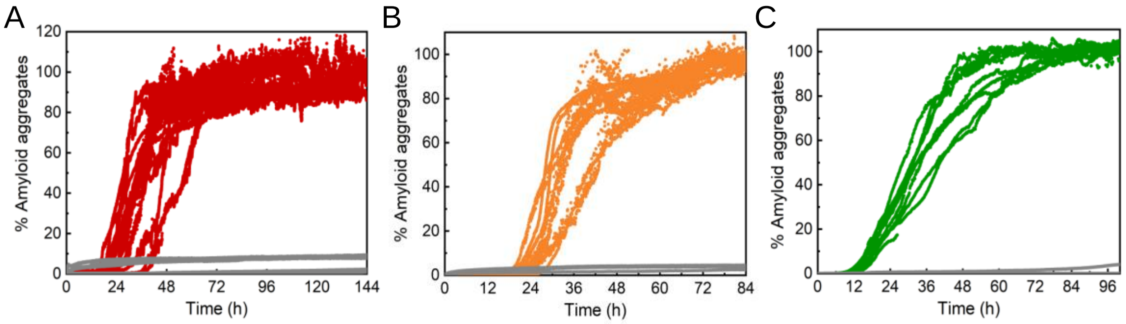


**Figure S10.** Removal of the active surfaces impairs αS aggregation. Aggregation kinetics of 100 µM αS in PBS pH 7.4 (37 °C) in the presence of 50 µM ThT and 10 % MeOH (A), 5 % TFE (B) and 2 M NaCl (C) in the presence of the air/water interface, without PMMA caps (colored curves) and without the interface, with PMMA caps (grey curves).

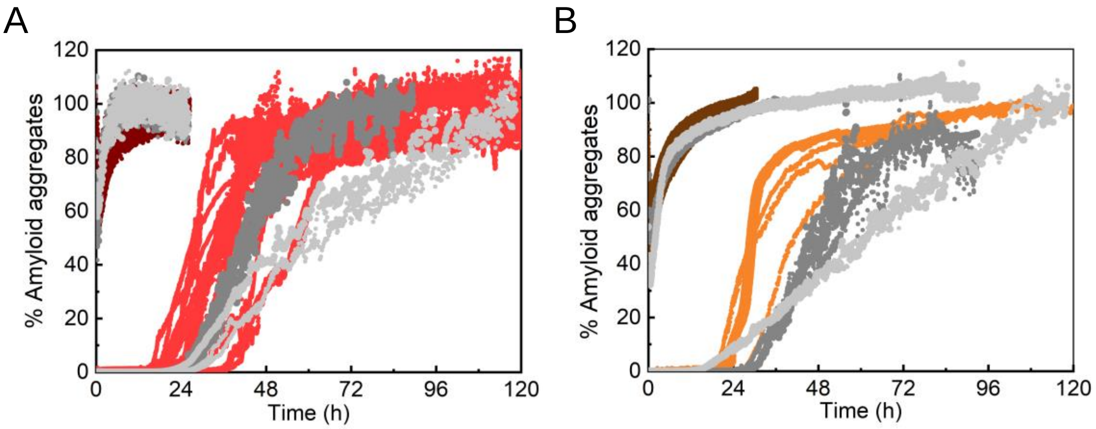


**Figure S11.** Comparison of the aggregational behavior of N-terminally or C-terminally modified αS variants. Aggregation kinetics of 100 µM αS in PBS pH 7.4 (37 °C) in the presence of 50 µM ThT and (A) 10 % or 35 % MeOH or (B) 5 % or 15 % TFE**.** The WT protein is shown in red (10 % MeOH) or dark red (35 % MeOH) in (A) and in orange (5 % TFE) or dark brown (15 % TFE) in (B), while the N-terminally acetylated protein is shown in dark grey and the C-terminally truncated (1-103) variant in light grey. In all cases, the modified protein variants behave similarly as the WT protein.

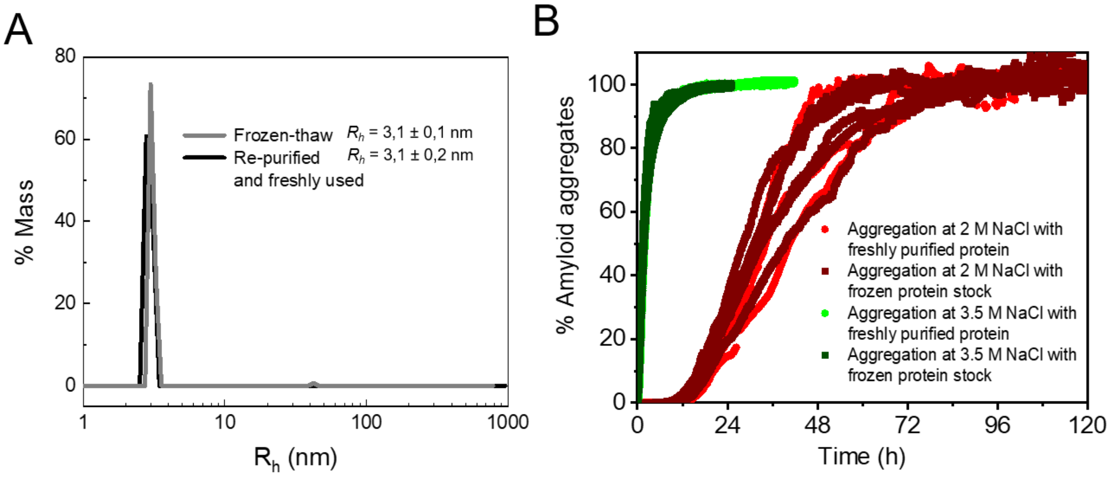


**Figure S12.** Assessment of the presence/absence of pre-formed nuclei in the protein batch samples. (A) Representative DLS profiles of the protein batch solutions before and after re-purification by SEC and analyzed/used immediately after column elution; grey and black lines, respectively. Most of the experiments presented in the manuscript were performed with protein batch aliquots that were stored at -80 °C just after purification by SEC (see Materials and Methods for a more detailed description of the purification protocol) and thawed only once when required. Some control experiments were also performed with protein batch samples that were re-purified by SEC and used immediately after protein elution from the column. When both types of protein samples were analysed by DLS for the presence of traces of pre-formed aggregates, both types of samples yielded virtually identical DLS profiles with only the monomeric protein, with a hydrodynamic radius of ~ 3.1 nm, apparently constituting 100% of the population. (B) Comparison of the aggregation kinetic profiles of the protein either with a protein batch solution purified by SEC and stored at -80 °C (aliquots thawed immediately before use) or with a protein batch solution that was re-purified by SEC and used immediately after column elution (dark and light colors, respectively). Aggregations through heterogeneous nucleation are shown in red symbols (PBS with 2.5 M NaCl) and aggregations through homogeneous nucleation in green symbols (PBS with 3.5 M NaCl).

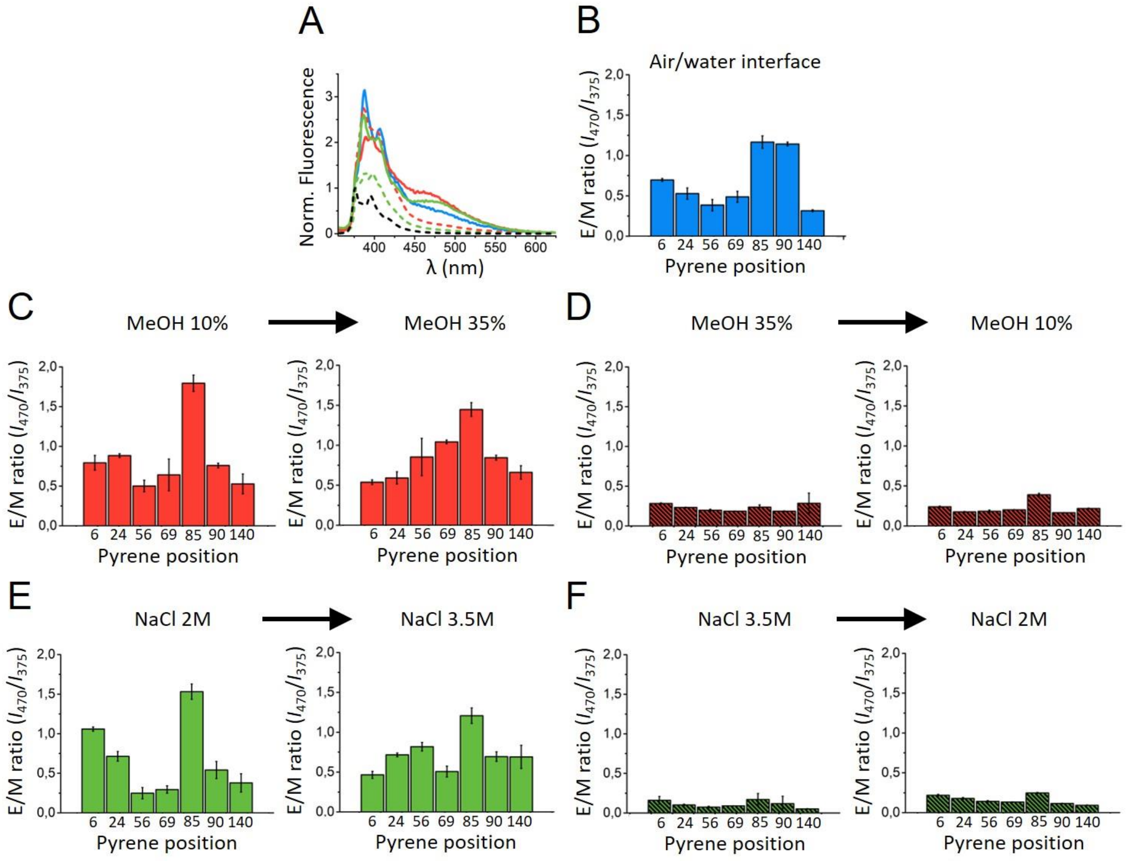


**Figure S13.** Intermolecular pyrene excimer formation analysis of the different β-sheet αS amyloid aggregates and structural stability of different αS aggregates in different environments. A) Representative pyrene fluorescence spectra of αS aggregates, labeled in position 24 of the protein primary sequence, formed at the air-water interface in PBS with shaking (blue), or in quiescence in PBS with 10 % MeOH (solid red line), or 2 M NaCl (solid green line), as well as aggregates formed in the bulk of the solutions in PBS with 35 % MeOH (dotted red line) or 3.5 M NaCl (dotted green line). B) Excimer/Monomer (E/M) ratio of pyrene fluorescence emission of the αS aggregates generated at the air-water interface in PBS with strong shaking. The pyrene moiety was added by cysteine maleimide reactions at different residue positions along the protein primary sequence (positions 6,24,56,69,85,90 and 140). C-F) Analysis of the pyrene fluorescence emission E/M ratio at different sequence positions of αS in the aggregates formed in the presence of 10 % MeOH (C), 35 % MeOH D), 2 M NaCl (E) and 3.5 M NaCl (F) before and after transferring them to other conditions as indicated by the arrows. Spectra were acquired at room temperature. In all cases, the aggregates maintained their original structural topology after being transferred to the new conditions, i.e., the parallel β-sheet aggregates remained parallel after being transferred to conditions where the antiparallel β-sheet topology was preferred (panels C and E), and *vice versa* (panels D and F). In the parallel β-sheet aggregates, which exhibited strong excimer formation, region-specific variations in E/M ratio were observed, indicating that the aggregates suffered a strong structural change at severe dehydrating conditions, in agreement with the remarkable loss of β-sheet signal observed by IR for the same experiment (see Figure 7 in the main text). It is also remarkable that the aggregates formed at the air/water interface by strong shaking or at quiescent conditions in the presence of 10 % MeOH or 2 M NaCl show significantly different E/M ratios for all the positions tested, indicating that the three types of parallel β-sheet aggregates have a different internal structure. However, in terms of stability, as indicated by the experiment showed in panels C and E, behave similarly. In the case of the aggregates formed in 35 % MeOH or 3.5 M NaCl, no significant E/M ratio was observed for any of the positions tested in αS, in agreement with their antiparallel β-sheet structure, which was maintained when they were transferred to increased protein hydration conditions, even if severely destabilized as observed by IR analysis (see Figure 7 in the main text).

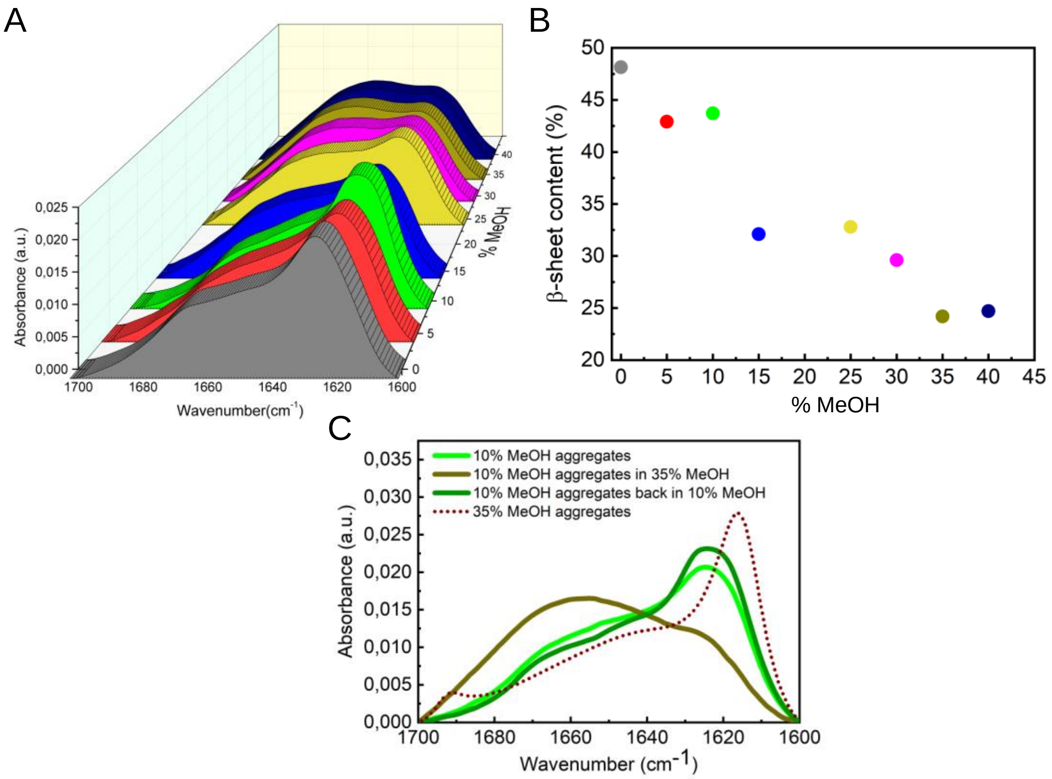


**Figure S14.** Analysis of the structural stability of the αS amyloid aggregates generated at 10 % MeOH when transferred to solutions with different MeOH concentrations. (A) IR spectra of aggregates at different MeOH concentrations in the 0-40 % range, under otherwise identical conditions (PBS, pH 7.4, 25 °C). The aggregates generated at 10 % MeOH were transferred to the different solutions and the analysis of their stability was performed after 15 h of incubation at room temperature under the new conditions. The aggregates did not dissociate when transferred at the different MeOH solutions (verified by SDS-PAGE analysis after ultracentrifugation; not shown), but significant structural changes in the aggregates were observed by IR analysis. (B) The β-sheet content of the aggregates under the different conditions was obtained from the quantitative analysis of the IR spectra (panel A). Upon increasing MeOH concentrations, there is a progressive decrease in β-sheet content, concomitant with an increase in likely random coil structure (the region of the spectra for random coil and α-helical structure largely overlaps in the amide I band [12]). The maximum β-sheet content was reached at 0 % MeOH, conditions at which the aggregates (originally generated at 10 % MeOH) show a very similar IR spectrum as the aggregates that were generated in PBS with shaking, suggesting that these two types of aggregates share similar structural features despite their different quaternary assembly (the aggregates generated in PBS under shaking exhibit a fibrillar morphology, while the aggregates generated in PBS + 10 % MeOH show a more globular-like morphology according to AFM imaging analysis; see Figures 1A and 1D in the main text). (C) Comparison of the IR spectra of the aggregates generated at 10 % MeOH when they remain at 10 % MeOH (green), when transferred to 35 % MeOH (brown) and when they are transferred back to 10 % MeOH (dark green). The spectrum of the aggregates generated at 35 % MeOH is also shown for comparison (dotted dark brown line). The structural destabilization of the 10 % MeOH aggregates at 35 % MeOH is reversed when the aggregates are transferred back to 10 % MeOH.

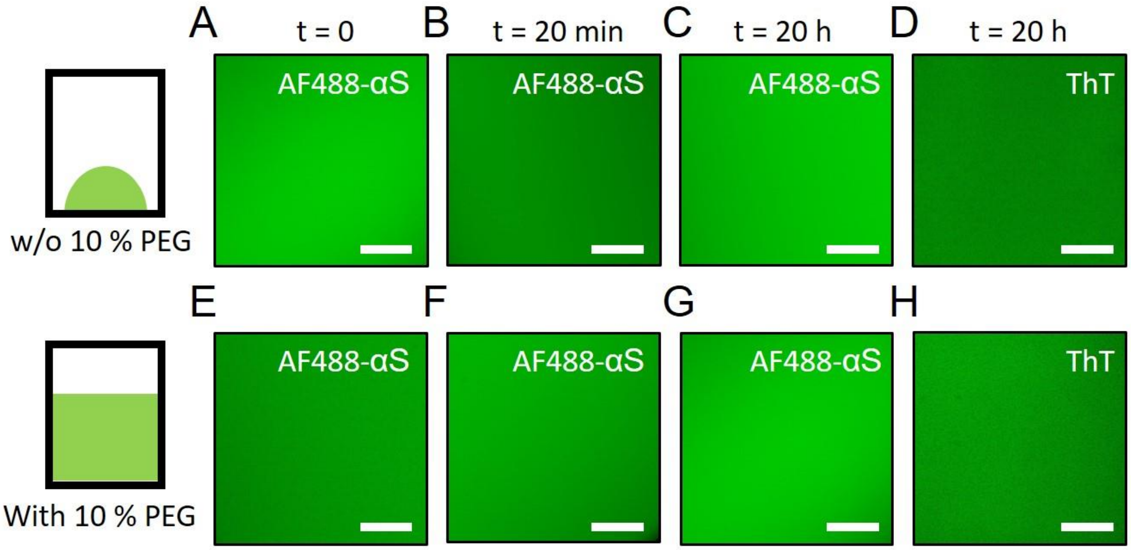


**Figure S15.** Analysis of the formation of αS droplets by LLPS. Representative widefield fluorescence microscopy images of protein samples containing 200 µM αS and 1 µM AF488-αS without PEG in a droplet setup (panels A-C) and with 10 % PEG in a microplate well (panels E-G). Panels A-C and E-G show the reaction at t = 0, t = 20 min and t = 20 h, respectively. Panels D and H are equivalent to panels C and G, respectively, with the exception that the sample contained only unlabeled αS and 100 µM thioflavin-T (ThT). Scale bar is 25 µm. A GFP excitation/emission filter set was used for all acquisitions. The images in the top panels show that 10% PEG is needed in order to trigger αS LPPS, and that, under those conditions (without PEG), no protein droplets or aggregates are, therefore, formed. In addition, the images in the bottom panels show that αS LLPS was not favored even after 20 h of incubation of protein solutions with 10 % PEG when the protein solutions were covering ca. 1/3 of the microplate wells, indicating that a droplet setup is needed for αS LLPS at the time scales analyzed in this study and that, therefore, the amyloid aggregate kinetics of αS in the presence of PEG ,described in the main text (section "Addition of macromolecular crowders also accelerates αS nucleation"), are triggered by heterogeneous nucleation at the air/water interface with the formation of parallel β-sheet amyloid aggregates. However, when the protein solutions were placed in a droplet setup at the surface of the wells of the microplates, LLPS was triggered and αS droplets were observed after 20-min incubation (see Fig. 8 in the main text). The aggregates generated inside the αS droplets by a liquid-to-solid transition (2 h for observing aggregation, 20 h for almost complete aggregation) showed a pyrene signature of intermolecular antiparallel β-sheet amyloid structure, thus suggesting that they were formed by homogeneous primary nucleation in the special microenvironment of the interior of the αS droplets, with a particularly low water activity in comparison with the solution conditions outside the protein droplets. DIC microscopy imaging of the samples yielded the same result, *i. e*., no LLPS nor protein aggregation was observed (not shown). Two independent triplicate experiments were performed.

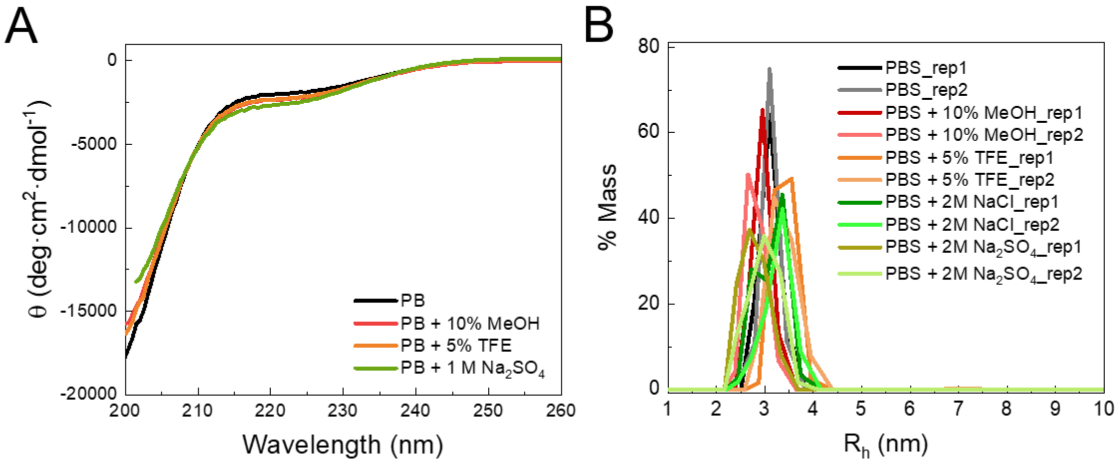


| Solution conditions | $R_h$ (nm) |
|---|---|
| PBS | 3,1 ± 0,1 |
| PBS + 10% MeOH | 2,9 ± 0,1 |
| PBS + 5% TFE | 3,4 ± 0,2 |
| PBS + 2 M NaCl | 3,2 ± 0,3 |
| PBS + 2 M $Na_2SO_4$ | 2,9 ± 0,3 |

**Figure S16.** Assessment of the general features of the structural ensemble of the monomeric protein in the presence of co-solvents and salts at concentrations that accelerate αS amyloid aggregation. (A) far-UV CD spectra. (B) DLS analysis. For the analysis of the secondary structure content by far-UV CD, the maximum salt concentration at which at least part of the spectrum could be recorded was 1 M $Na_2SO_4$.

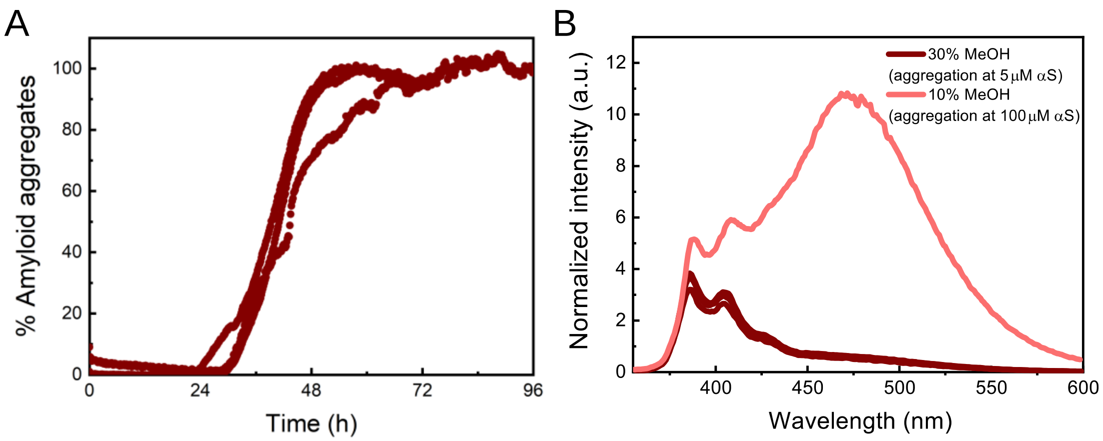


**Figure S17.** αS aggregation at 5 µM protein concentration in PBS pH 7.4, 30 % MeOH. A) Aggregation kinetics of 5 µM αS in PBS pH 7.4 (37 °C) in the presence of 30 % MeOH with 50 µM ThT. (B) Fluorescence spectra of 5 µM pyrene-labelled αS aggregates formed in the presence of 10 % and 30 % MeOH. Only labelled αS was used, carrying the pyrene moiety in position 85. Spectra were normalised to $I_{375}$. The data indicate that αS is able to aggregate at low protein concentrations, in the very low micromolar range, under conditions of homogeneous nucleation with the formation of antiparallel β-sheet aggregates.

### Supporting Tables

**Table S1.** Summary of the main features of αS aggregation under the different conditions used in this study

| Condition[1] | Aggregation kinetics | | Aggregates | | |
|---|---|---|---|---|---|
| | Duration of lag phase | Apparent nucleation type | β-sheet arregement | % β-sheet (N)[2] | Stability in PBS |
| **PBS a/w interface[3]** | Extremely slow (>200h) | Heterogeneous | Parallel | 54 ± 4 (9) | Yes |
| **+ PTFE Bead** | Slow (ca. 10h) | Heterogeneous | Parallel | 51 ± 4 (2) | Yes |
| **+ SUVs[4]** | Slow (ca. 10h) | Heterogeneous | Parallel | 46 ± 3 (5) | Yes |
| **+ 5% TFE** | Slow (ca. 20h) | Heterogeneous | Parallel | 49 ± 3 (3) | Yes |
| **+ 6% MPD** | Slow (ca. 30-40 h) | Heterogeneous | Parallel | 42 ± 3 (4) | Yes |
| **+ 7% i-PropOH** | Slow (ca. 50 h) | Heterogeneous | Parallel | 50 ± 8 (3) | Yes |
| **+ 10% MeOH** | Slow (ca. 30-40 h) | Heterogeneous | Parallel | 41 ± 9 (7) | Yes |
| **+ 15% TFE** | Fast (<5-10 min) | Homogeneous | Antiparallel | 60.2 ± 0.6 (8) | No |
| **+ 24% MPD** | Fast (<5-10 min) | Homogeneous | Antiparallel | 59 ± 2 (2) | No |
| **+ 26% i-PropOH** | Fast (<5-10 min) | Homogeneous | Antiparallel | 58 (1) | No |
| **+35% MeOH** | Fast (<5-10 min) | Homogeneous | Antiparallel | 57 ± 6 (2) | No |
| **+ 2M NaCl** | Slow (ca. 20 h) | Heterogeneous | Parallel | 43 (1) | Yes |
| **+ 2M $Na_2SO_4$** | Slow (ca. 5 h) | Heterogeneous | Parallel | 51.44 (1) | Yes |
| **+ 3.5M NaCl** | Fast (<5-10 min) | Homogeneous | Antiparallel | 20.0 ± 0.1 (2) | No |
| **+ 3M $Na_2SO_4$** | Fast (<5-10 min) | Homogeneous | Antiparallel | 20 (1) | No |
| **+ 150 g/l Dext 70** | Slow (ca. 10 h) | Heterogeneous | Parallel | 42 ± 4 (3) | Yes |
| **+ 150 g/l Ficoll 70** | Slow (ca. 10 h) | Heterogeneous | Parallel | 45 ± 1 (2) | Yes |
| **+ 150 g/l PEG 8** | Slow (ca. 5-10 h) | Heterogeneous | Parallel | 40 ± 9 (2) | Yes |
| **+ 100 g/l PEG8 (LLPS)** | Fast | Homogeneous[5] | Antiparallel | - | - |
| **Toxic Oligomer[6]** | - | - | Antiparallel | 35 ± 5 | Yes |

[1] All conditions contained 100 µM αS in PBS pH 7.4 and aggregations were performed at 37 °C in quiescence, except for the conditions referred to "PBS a/w interface", "+ SUVs" and "+100 g/L PEG8 (LLPS)". In the latter, 200 µM αS in PBS pH 7.4 was used. The concentration of salts is given in ionic strength. [2] *N* refers to the number of replicas used in the analysis; the numbers represent the average and standard deviation. [3] *a/w interface* stands for air/water interface. Under this condition no aggregation is observed for 100 µM µS and in quiescent conditions (lag phase longer than 250 h), so the aggregates were generated with 500 µM αS and shaking. [4] The solution condition in this case was phosphate buffer pH 6.5, 37 °C in quiescence. [e] Data on the toxic αS oligomer was obtained previously [13–16] and included here for comparison. [5] The apparent type of primary nucleation was not directly determined but the different experimental evidences point to a homogeneous nucleation (see main text of the article). [6] This type of oligomer has an intermediate structure between the monomers and the fully-formed fibrils and is highly toxic to cells. We recently demonstrated that they are formed during the protein lyophilization process[13].

### Author Contributions

NC design the study and JDC performed most of the experiments under the guidance of SWC, JS, FMG, IA, JLRA, CMD and NC. PG performed all the pyrene-based and LLPS experiments. VS generated the microfluidic system. JDC, PG and NC analyzed the data and wrote the paper. All authors have given approval to the final version of the manuscript.
